# Horizontally acquired mitochondrial alternative oxidase contributes to springtail bioenergetics and life belowground

**DOI:** 10.64898/2026.06.01.729305

**Authors:** Ryan J. Weaver

**Author notes:** Ryan J. Weaver, **Email:**.

## Abstract

Springtails are tiny hexapods that occupy habitats ranging from above the soil surface to hypoxia-prone belowground environments, yet the physiological mechanisms enabling this ecological expansion have remained unresolved. Here, I take an evolutionary bioenergetics approach to show that a mitochondrial alternative oxidase (AOX) acquired by horizontal gene transfer in the springtail ancestor became functionally integrated into animal physiology and was preferentially retained in lineages occupying low-oxygen habitats. Surveying 202 springtail genome assemblies, I identified 65 high-confidence AOX loci in 48 species, each embedded within otherwise typical springtail genomic neighborhoods. Phylogenetic and motif-based analyses support an oomycete donor and indicate ancestral acquisition followed by repeated loss, especially in aboveground taxa. High-resolution respirometry and hypoxia-exposure experiments further show that AOX is active only in AOX-positive species and is associated with hypoxia tolerance. These results identify horizontal gene transfer as a source of ecophysiological innovation in animals and suggest that acquired mitochondrial functions can help shape ecological sorting across environmental gradients, with implications for soil ecosystem processes.

**Significance Statement:** Horizontal gene transfer can introduce new genes into animal genomes, but few cases connect gene origin to physiology and ecology. Springtails are abundant soil mesofauna that help define ecological systems and nutrient flux yet physiological challenges that subterranean environments impose raise questions regarding the mechanisms of adaptation to life below ground. This study shows that springtails acquired mitochondrial alternative oxidase (AOX), a respiratory enzyme widely associated with stress tolerance in plants, fungi, protists, and some animals, from an oomycete donor early in their evolutionary history and that AOX has been retained in hypoxia-tolerant species and incorporated it into the physiology as a potential mechanism for tolerance of belowground conditions. These findings link macroevolutionary patterns to bioenergetic processes that help shape ecological distribution and tolerance of environmental stressors, with implications for biodiversity and ecosystem function.

## Introduction

Belowground environments impose strong selective pressures on animal physiology. Compared to aboveground habitats, soils experience low and fluctuating oxygen availability due to microbial respiration, soil structure, moisture, and diffusion limitation. Species that habitually occupy these environments often show traits associated with life below ground, including reduced pigmentation, eye reduction or loss, and morphological changes that may facilitate movement through the soil matrix (1–5). From a bioenergetic perspective, one of the most consequential features of belowground habitats is low and dynamic oxygen availability (6–10). This is especially important because mitochondria are the principal consumers of oxygen in the cell. In addition to synthesizing ATP through oxidative phosphorylation (OXPHOS), mitochondria play essential roles in central carbon metabolism and in the production or regulation of key cellular components, including tricarboxylic acid intermediates, iron–sulfur clusters, heme, and reactive oxygen species (ROS). Many of these functions depend directly or indirectly on the continued oxidation of reducing equivalents and, in aerobic cells, on the availability of oxygen as the terminal electron acceptor. When oxygen becomes limiting, mitochondria must be reprogrammed to avoid the damaging consequences of respiratory stress (9, 11–15).

Mitochondrial respiration in animals is often conceptualized as a linear electron transport chain in which electrons enter through complex I or complex II, pass through coenzyme Q, complex III, cytochrome c, and finally complex IV, where oxygen is reduced to water (Fig. 1). This chain-like framework is useful, but it is not representative of the broader diversity of mitochondrial respiratory architecture across eukaryotes (16). In many lineages, respiration is better portrayed as a branched electron transfer system, where multiple electron entry points converge at the coenzyme Q pool and more than one terminal oxidase can accept electrons and catalyze the reduction of oxygen to water (17, 18). This branched architecture provides metabolic flexibility, which is especially relevant for understanding how mitochondria function under environmental or cellular stress (19–23).

**Figure 1.**
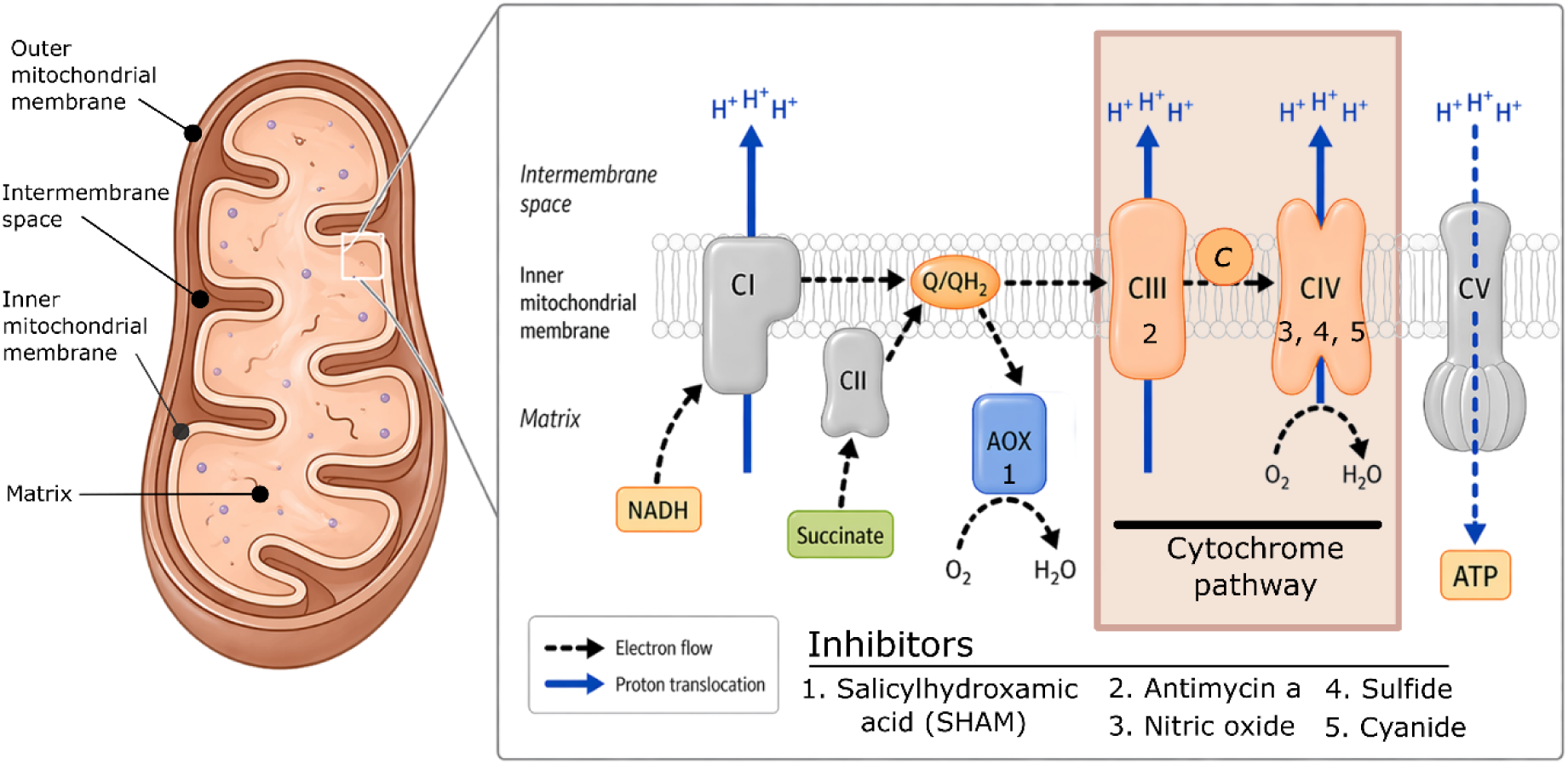
Schematic view of the mitochondrial electron transport system highlighting the alternative oxidase (AOX) and cytochrome pathways. Electrons from NADH and succinate enter the ubiquinone pool through complexes I and II, respectively. From reduced ubiquinone (QH₂), electrons can flow through the proton-translocating cytochrome pathway via complex III, cytochrome c, and complex IV, supporting ATP production by complex V, or through AOX, which reduces O₂ to H₂O while bypassing complexes III and IV. Dashed black arrows indicate electron flow, blue arrows indicate proton movement, and numbered labels mark representative inhibitors of AOX or cytochrome-pathway components.

A key component of this branched architecture is alternative oxidase (AOX), a nuclear-encoded diiron terminal oxidase that transfers electrons from the reduced coenzyme Q pool directly to oxygen, bypassing complexes III and IV (Fig. 1)(24, 25). On a per electron basis, AOX is less efficient at ATP synthesis compared to the cytochrome pathway because it bypasses proton-translocating steps of complexes III and IV. Yet this apparent inefficiency may provide an important physiological advantage: metabolic flexibility. When electron flux through the cytochrome pathway is constrained, AOX can preserve electron flow, limit over-reduction of the electron transfer system, and help maintain redox balance (21, 22, 26–31). Under hypoxia, and especially during reoxygenation, these effects may be particularly important because restricted cytochrome-pathway flux can result excessive reactive oxygen species production, and oxidative damage (9, 32–35).

AOX and other alternative respiratory enzymes are widespread across eukaryotes but patchily distributed among animals (18, 36). AOX has been lost in several metazoan lineages, including vertebrates and most arthropods. Recent work, however, has suggested that AOX has been regained in several animal groups, putatively through horizontal gene transfer (18). Using sequence motif analysis, phylogenetics, and BLAST searches, Weaver and McDonald (18) proposed that AOX was horizontally acquired in rotifers, nematodes, fungus gnats, and springtails, with putative donor lineages including fungi in nematodes and rotifers and protists in fungus gnats and springtails. These findings create an opportunity to address a fundamental evolutionary question: how do environmental conditions and natural selection interact to transform horizontally transferred genetic material into functionally integrated and ecologically relevant components of animal physiology?

Springtails (Hexapoda; Collembola) are a taxonomically and ecologically diverse group of soil-associated hexapods and provide an especially useful system in which to address this question. Across multiple springtail lineages, species occupy habitats spanning an aboveground to belowground gradient and are categorized into three broad “life forms” (37). Epedaphic species primarily inhabit the soil surface and leaf litter, hemiedaphic species occupy the lower litter layer and upper soil horizon, and euedaphic species live deeper within the soil matrix (38, 39).

Because these ecological categories differ in expected exposure to oxygen limitation (8, 40, 41), springtails represent a natural comparative framework to test whether AOX is associated with adaptation to hypoxia-prone environments. If AOX maintains core mitochondrial functions during oxygen stress, its retention may be favored in lineages that more frequently encounter low-oxygen microhabitats.

Here, I provide evidence that AOX in springtails represents a genuine case of horizontal gene transfer followed by functional integration into host physiology as an adaptation to belowground life. First, I examine genomic signatures of springtail AOX to distinguish true integration from an artifact of contamination. If AOX is genuinely incorporated into springtail genomes, then AOX-positive species should possess orthologous sequences embedded in native springtail genomic context and show consistent motif and phylogenetic signatures of protist origin, rather than patterns expected of bacterial, fungal, or other environmental contaminants. Second, I use high-resolution respirometry to determine whether AOX contributes to mitochondrial function in springtail species that possess it. If AOX is functionally integrated, then respiration in AOX-positive species should show partial resistance to inhibition of the cytochrome pathway and sensitivity to AOX inhibition, whereas AOX-negative species should lack respiratory sensitivity to AOX inhibitors and remain fully dependent on the cytochrome pathway. I also perform hypoxia-exposure experiments to test a prediction of an AOX-hypoxia hypothesis: AOX-positive species tolerate hypoxia exposure whereas AOX-negative species do not. Finally, I use comparative phylogenetic analyses to reconstruct the evolutionary history of AOX and another hypoxia-relevant protein (hemocyanin (42)) in springtails and test whether their retention is associated with ecological transitions into hypoxia-prone habitats. If AOX contributes to coping with hypoxia, I predict that it will be preferentially retained in hemiedaphic and euedaphic lineages and more frequently lost in epedaphic species.

## Results

### AOX has been integrated into the nuclear genome of Folsomia candida

In the *Folsomia candida* RefSeq genome assembly, AOX resides within a 76-gene genomic region in which every non-AOX gene returned a top BLAST hit to a metazoan sequence, whereas AOX returned top hits to oomycete protists (Fig. 2c). The AOX gene model consists of four exons and three introns with splice-site motifs typical of eukaryotic genes and the expected N-terminal mitochondrial targeting signal (Fig. S1C, Table S4, S5). The coding region contains all four catalytic iron-binding sites required for AOX function (Fig. 2b). The AOX scaffold has median sequencing depth and coverage typical of the broader genome assembly (Fig. S1C). Pairwise alignments near these iron-binding sites showed conservation between *F. candida* and the oomycete protist *Pythium aphanidermatum* to the exclusion of typical metazoan AOX motifs (Fig. 2a). No rRNA features were detected on the AOX-containing scaffold (SI Appendix).

**Figure 2.**
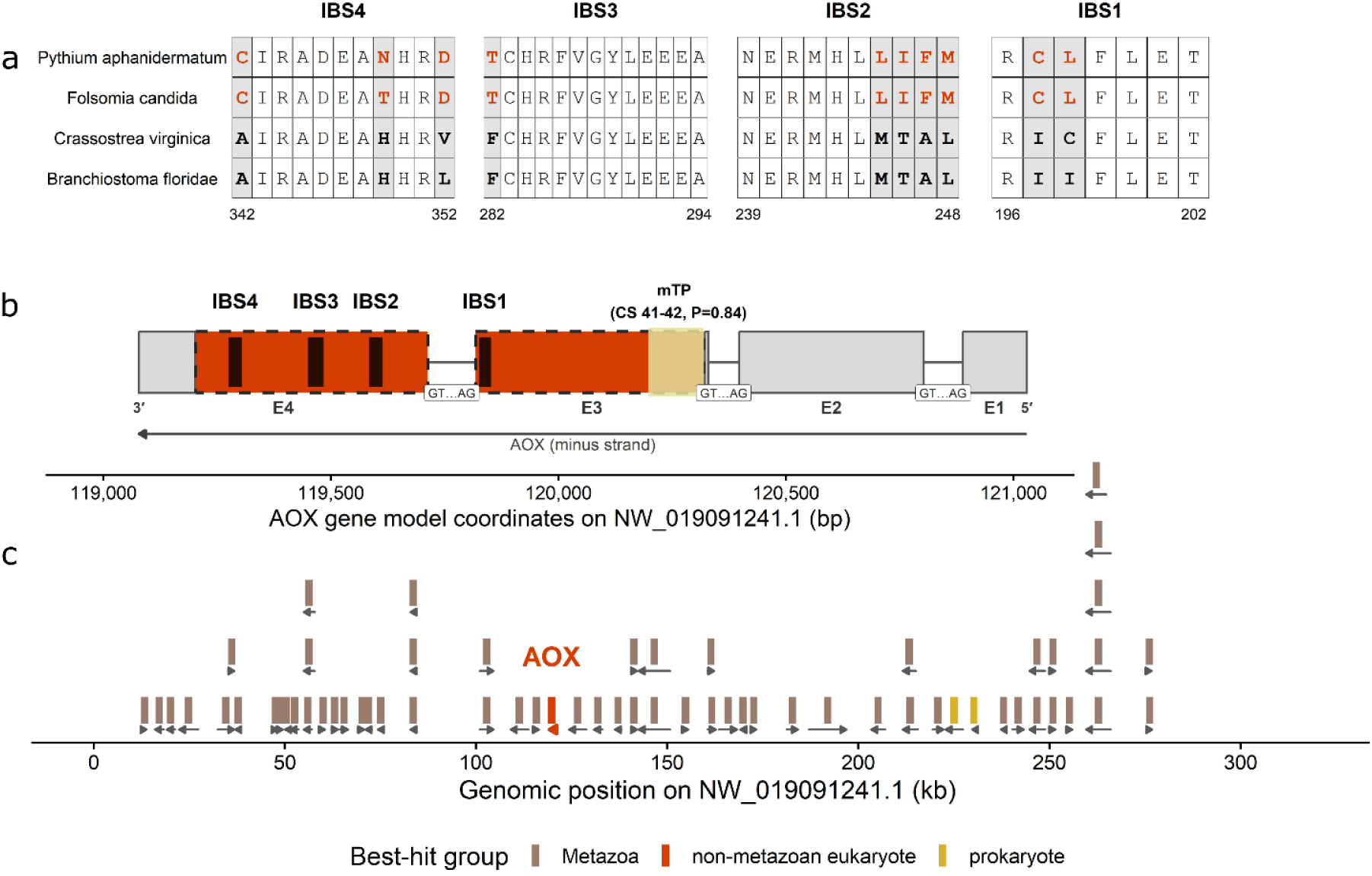
The genomic context of AOX in *Folsomia candida*. a) Multiple sequence alignment of the four requisite iron binding sites (IBS) of AOX amino acid sequences from a protist (*Pythium aphanidermatum),* a springtail (*Folsomia candida)*, and two other metazoans: a bivalve (*Crassostrea virginica*) and a lancelet (*Branchiostoma floridae*). Shaded positions highlight where springtail and protist amino acids are conserved to the exclusion of the other metazoans. b) the AOX gene model from the RefSeq genome assembly showing four exons (gray, UTR; red, coding sequence). The mitochondrial targeting peptide region and cleavage site are highlighted in yellow. Dark bars show the position of the four iron-binding sites (IBS). c) BLAST results for genes within a 200 kb window of the AOX scaffold. Brown bars show genes most similar to other metazoans, red shows non-metazoan eukaryotic hits, yellow shows prokaryotic hits.

### AOX microsyteny

To test whether the RefSeq AOX locus is embedded within a conserved *F. candida* genomic neighborhood, I projected the 16 protein-coding genes flanking AOX in the RefSeq assembly onto three additional *F. candida* assemblies. In all three assemblies, AOX was successfully anchored and all 16 non-AOX flanking genes were recovered on the expected side of AOX with complete concordance across flanking positions (Fig. S2, Table S6). This conserved local microsynteny strongly supports interpretation of the AOX locus as a genuine component of the *F. candida* nuclear genome.

### AOX presence and genomic context in publicly available springtail genomes

Mining 202 springtail genome assemblies recovered 75 AOX-positive loci. Because some species were represented by more than one genome assembly, these candidates were treated as accession-level records rather than independent species-level observations. Candidate AOX sequence support varied among accessions, reflecting differences in assembly fragmentation, AOX model completeness, and local scaffold context (Fig. S3). Nevertheless, many candidates were supported by stitched tBLASTN reconstructions that recovered three or four conserved AOX iron-binding-site motifs, consistent with bona fide AOX coding sequence rather than short nonspecific similarity.

Fragment-aware inspection of AOX-positive loci showed that the recovered candidates differed substantially in local structure (Fig. S4). Some accessions contained compact AOX-like intervals with strong stitched-candidate support and local sequence windows fully within longer contigs. Others were positioned near contig boundaries or were reconstructed from multiple AOX-like fragments, consistent with the fragmented nature of many available springtail assemblies. Thus, the genomic context analysis provided local scaffold inspection and locus-quality assessment, but it was not consistently strong enough across accessions to recover exact flanking-genes.

Neighborhood projection of AOX loci further supported the interpretation that many AOX-positive candidates occur within broader gene-containing regions of the genome. Of 75 AOX-positive assemblies, 27 retained projected intervals (i.e., gene-like regions) on both sides of AOX, including *Oncopodura yosiiana* (4.3 kb; 2.1 kb), *Thalassaphorura encarpata* (7.1 kb; 3.7 kb), and *Podura aquatica* (8.2 kb; 5.4 kb). An additional 22 accessions had recoverable intervals on only one side of AOX, whereas 26 lacked immediately recoverable flanking gene-like intervals on either side. Thus, 49 of 75 AOX-positive windows had at least one nearby projected interval, but local context varied substantially among assemblies. Taken together, these results indicate heterogeneous local structure across AOX-positive loci, while also showing that many candidates are not isolated AOX-like sequence islands and instead occur within broader genic regions of springtail assemblies.

### AOX phylogeny and donor placement

In the broad AOX phylogeny, springtail AOXs formed a monophyletic clade outside canonical metazoan AOXs and instead occupied an oomycete-associated region of tree space (Fig. 3a,b). Approximately unbiased topology tests supported this placement. The unconstrained reduced maximum-likelihood tree and the topology forcing springtail AOX as sister to sampled oomycete AOXs were nearly identical in fit (Delta lnL = 0.09, p-AU = 0.54; Fig. 3c), whereas forcing springtails to be sister to metazoan AOXs produced a significantly worse topology (Delta lnL = 18.19, p-AU = 0.008). These results support an oomycete-associated origin of springtail AOX and reject vertical inheritance from canonical metazoan AOX.

**Figure 3.**
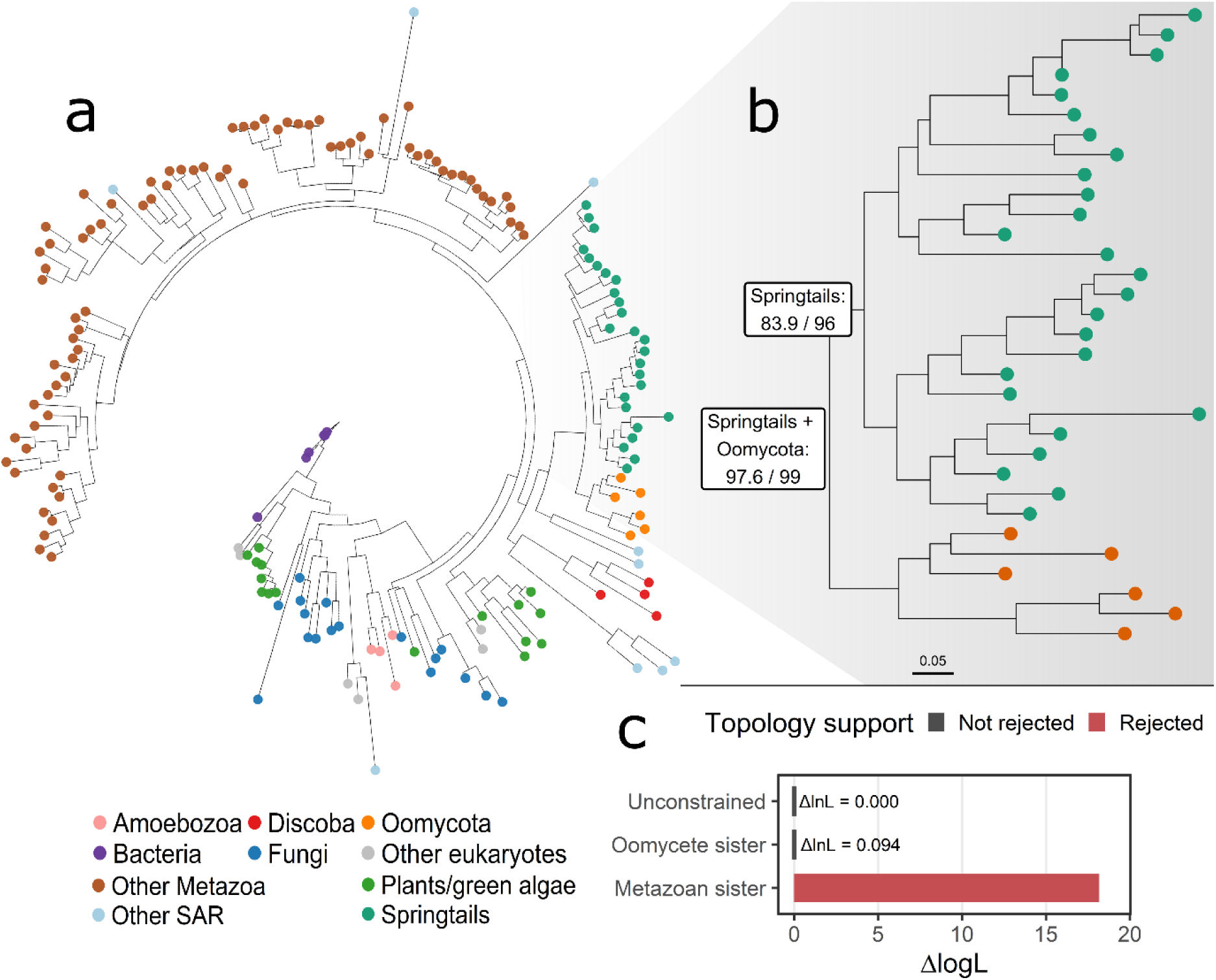
AOX phylogeny and topology-test support for springtail AOX placement. (a) Maximum-likelihood AOX protein phylogeny inferred from a 165-amino-acid alignment of 172 sequences. (b) Enlarged view of the springtail-containing region of the tree. Springtail AOX forms a monophyletic clade adjacent to sampled oomycete AOXs. Node labels are SH-aLRT/UFBoot values; support for the springtail + oomycota neighboring node is 97.6/99, and support for the springtail clade is 83.9/96. Scale bar indicates amino acid substitutions per site. (c) Approximately unbiased topology tests on a reduced, manually curated AOX dataset. The unconstrained reduced ML tree and the oomycete-sister topology were retained (ΔlnL = 0.09, p-AU = 0.54), whereas the metazoan-sister topology was rejected (ΔlnL = 18.19, p-AU = 0.008).

### Functional validation of AOX in Springtails

High-resolution respirometry confirmed a functional AOX pathway in *F. candida* and *Proisotoma minuta*, which are AOX-positive euedaphic and hemiedaphic species, respectively. After cytochrome-pathway inhibition, oxygen consumption persisted at approximately 16% of maximal respiration and was abolished only after addition of the AOX inhibitor SHAM (*F. candida*: (mean ± s.e) = 146.5 ± 6.9 pmol O_2_ min ^-1^ mg ^-1^, n = 5; *P.minuta*: 146.1 ± 12.8, n = 5, Figure 4a). In contrast, mitochondrial respiration in AOX-negative species was abolished by cytochrome-pathway inhibition alone. Adding cytochrome-pathway inhibitor at the onset of respirometry reinforced this pattern. Only AOX-positive species maintained cytochrome pathway inhibitor-resistant respiration, while AOX inhibition alone did not abolish mitochondrial respiration in any species (Fig. 4b).

**Figure 4.**
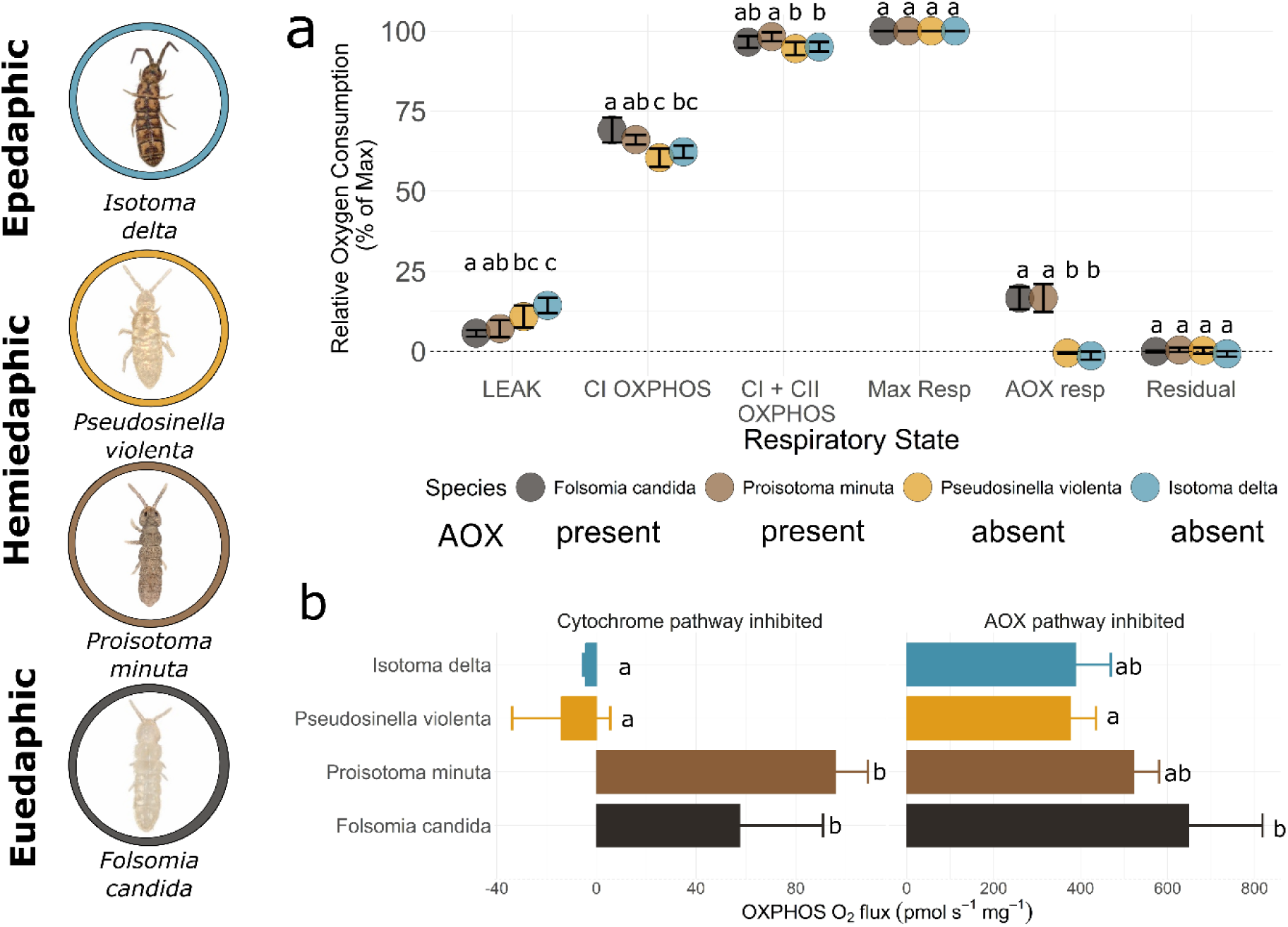
Functional validation of AOX in AOX-positive species. Respiration rates measured in AOX-positive and AOX-negative springtail species. a) Relative oxygen fluxes (% of maximum respiration rate) were determined for each respiratory state by sequential addition of pyruvate + malate (LEAK), ADP (CI OXPHOS), succinate (CI + CII OXPHOS), CCCP (Max Resp), antimycin a (cytochrome inhibitor; AOX resp), and salicylhydroxamic acid (AOX inhibitor; Residual). Circles and error bars show mean ± s.e., n = 5. b) CI + CII OXPHOS oxygen consumption rates measured when cytochrome pathway inhibitor or AOX inhibitor was added at the beginning of the respirometry measurements. Bars and error bars show means ± s.e., n = 3. Means that do not share a letter are statistically significant at α = 0.05.

### AOX-positive species tolerate hypoxia, AOX-negative species do not

After hypoxia and 24 h reoxygenation, the two AOX-positive species showed consistently high recovery (Fig. 5). *F. candida* showed recovery in 41/42 individuals after 4 h hypoxia and 94/95 after 16 h hypoxia, and *P. minuta* recovered in 45/48 and 87/94 individuals, respectively. In contrast, AOX-negative *Pseudosinella violenta* recovered in 23/40 individuals after 4 h hypoxia but only 10/96 after 16 h hypoxia, whereas AOX-negative *Isotoma delta* failed to recover after either exposure duration. Species effects were significant within both hypoxia treatments by Fisher’s exact tests (4 h and 16 h: p < 2.2 x 10^-16^) and by Firth logistic regression (4 h: likelihood-ratio test = 93.26, df = 3, p < 0.001, n = 154; 16 h: likelihood-ratio test = 378.94, df = 3, p < 0.001, n = 371). All species recovered completely under normoxic chamber controls (48/48 per species), indicating that handling and chamber exposure alone did not explain the hypoxia response.

**Figure 5.**
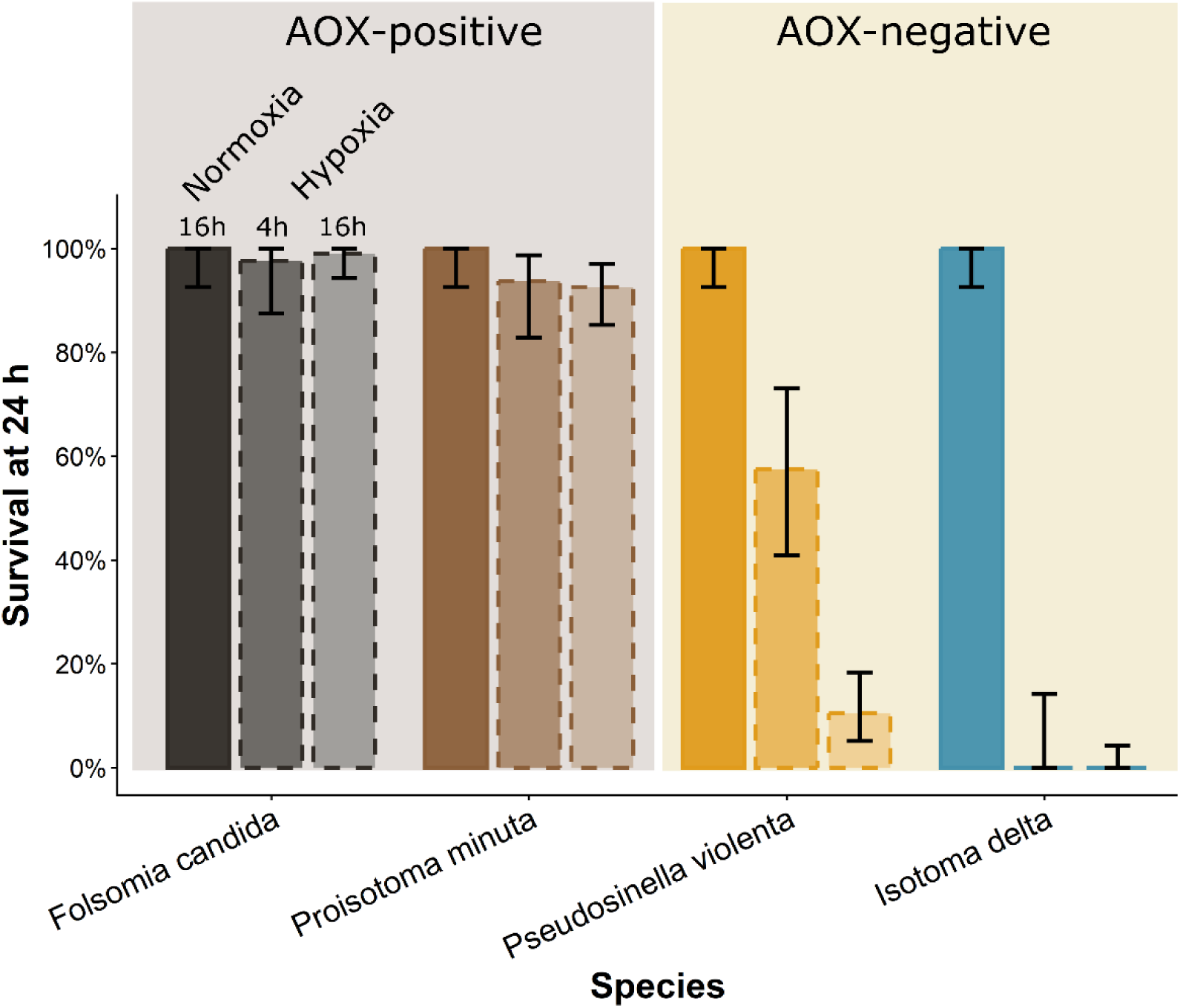
AOX-positive springtails maintain survival after hypoxia, whereas AOX-negative species show reduced or no survival. Proportion of individuals displaying normal activity 24 h after reoxygenation following 16 h normoxia, 4 h hypoxia, or 16 h hypoxia. Bars show exact binomial proportions; error bars indicate 95% confidence intervals. Bars are colored by species, and exposure groups are distinguished by fill opacity and line type.

### AOX evolution in springtails

Ancestral-state reconstruction supported AOX absence at the deeper hexapod root but high posterior probability of AOX presence at most recent common ancestor of springtails (Fig. 6a). Under the equal-rates model with a fixed AOX-absent hexapod root, the posterior probability of AOX presence at the springtail node was 0.98 (logLik = -15.7), and an unconstrained-root analysis gave a similar result (present = 0.96, logLik = -15.7). Sensitivity analyses using soft root priors and all-rates-different models were consistent with ancestral acquisition along the springtail stem followed by repeated losses, primarily in aboveground lineages (SI Appendix).

**Figure 6.**
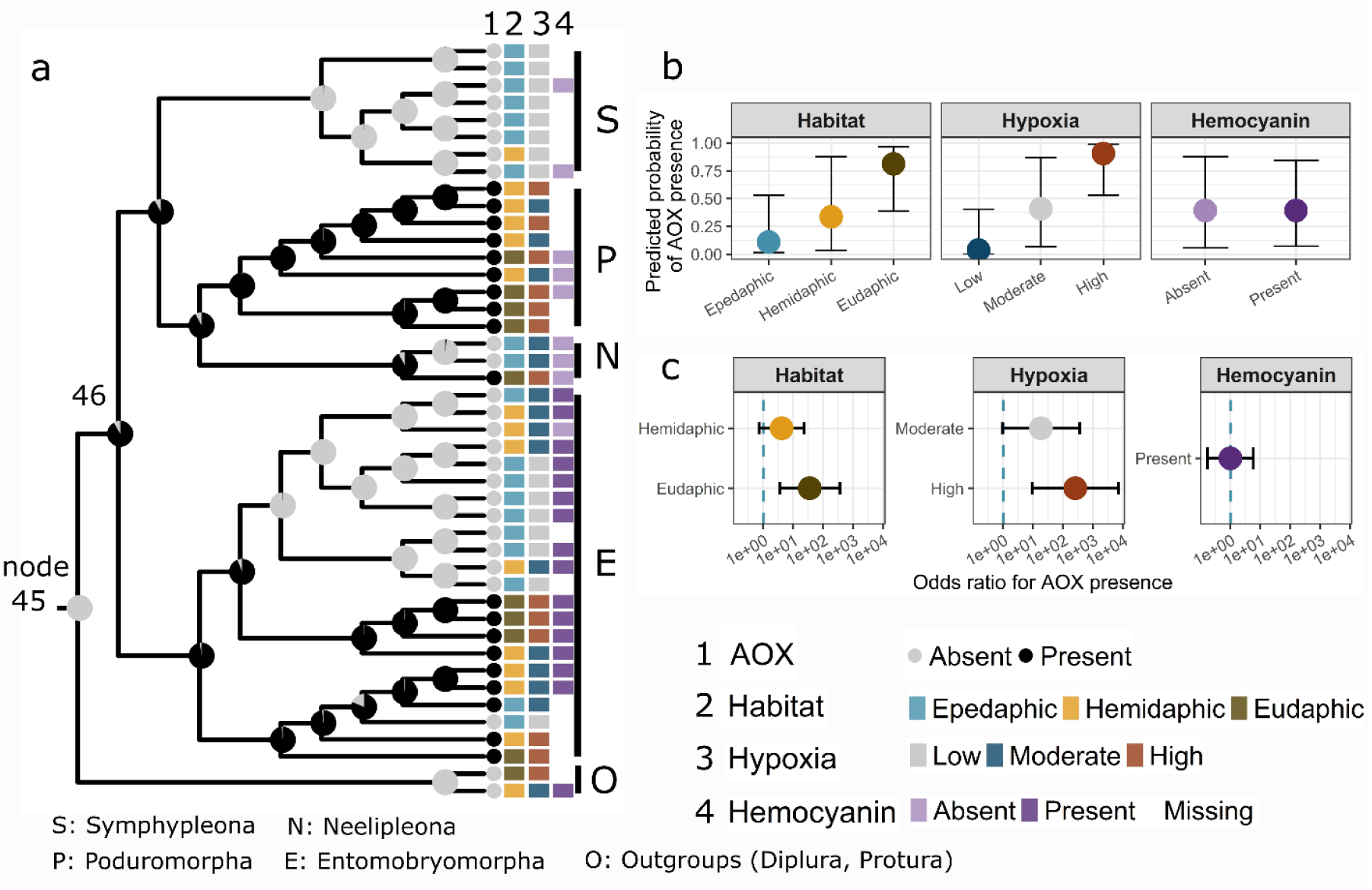
Comparative phylogenetics of AOX, hemocyanin, habitat, and hypoxia risk in springtails. a) Ancestral state reconstruction of AOX in springtails. Gray circles at the tips represent AOX-negative species, black circles show AOX-positive species. Pie charts at the internal nodes up to the MRCA of springtails represent the likelihood of AOX presence (black) or absence (gray) respectively. Colored bars at each tip map habitat, hypoxia risk, and hemocyanin absence/presence. b) Predicted probability of AOX presence from phylogenetic logistic regression models for habitat type, hypoxia risk, and hemocyanin presence. Circles and error bars represent the mean probability and 95% confidence limits. c) Predicted odds ratios of AOX presence relative to the epedaphic, low hypoxia, or hemocyanin-negative states.

Pagel’s discrete tests supported correlated evolution between AOX presence and high hypoxia exposure (Table S17; logL_ind = -35.79, logL_dep = -28.21, LR = 15.16, p = 0.00051; AIC_ind = 75.58, AIC_dep = 64.42) and between AOX presence and euedaphic occupancy (logL_ind = - 32.95, logL_dep = -27.45, LR = 10.99, p = 0.004; AIC_ind = 69.9, AIC_dep = 62.9). Phylogenetic logistic regression produced a concordant result: high-hypoxia species had significantly greater odds of possessing AOX than low-hypoxia species (Table S16; β = 5.54 ± 1.68 SE, z = 3.30, p = 0.001), and euedaphic species had significantly greater odds of possessing AOX than epedaphic species (β = 3.57 ± 1.17 SE, z = 3.03, p = 0.002). Moderate-hypoxia and hemiedaphic categories showed weaker effects with confidence intervals that included 1 (Fig. 6 b,c, Table S16). In contrast, hemocyanin presence was not significantly associated with hypoxia exposure, habitat, or AOX presence (Fig. 6 b,c, Table S17). Overall, AOX retention, but not hemocyanin presence, was associated with belowground life and hypoxia risk in sampled springtails.

## Discussion

Here, I show that mitochondrial alternative oxidase (AOX) in springtails is a horizontally acquired respiratory gene that has been incorporated into animal physiology and preferentially retained in lineages occupying hypoxia-prone environments. This study establishes an evolutionary bioenergetics framework for understanding how horizontally acquired metabolic genes can become functionally integrated into animal physiology and shaped by ecological selection. Specifically, it connects three levels of evidence that are often considered separately: genomic evidence for a non-metazoan origin, physiological evidence that the acquired gene contributes to mitochondrial respiration, and comparative evidence that AOX retention is associated with ecological exposure to hypoxia. Together, these results support a model in which a foreign genetic element became part of the springtail bioenergetic toolkit and subsequently experienced ecological sorting across the aboveground-belowground habitat gradient.

Claims of horizontal gene transfer in soil animals require special care because soil-associated genomes, cultures, and sequencing libraries are exposed to bacterial, fungal, protist, and endosymbiont DNA. Sequence similarity alone cannot discern between contaminant and genuine horizontal transfer of DNA. In this study, several lines of evidence support the conclusion that springtail AOX is a genuine nuclear gene of non-metazoan origin rather than a contaminant. AOX occurs in typical host genomic context, is surrounded by genes with metazoan affinities, and is recovered in conserved microsyntenic neighborhoods across *Folsomia candida* assemblies.

The gene also has eukaryotic exon-intron structure and contains the conserved catalytic iron-binding residues required for AOX function. At the same time, springtail AOX lacks the motif signatures expected of vertically inherited animal AOX and instead carries oomycete-like iron-binding-site motifs. Phylogenetic analyses place springtail AOXs outside canonical metazoan AOXs, and topology tests reject a metazoan-sister placement while retaining an oomycete-sister topology. Therefore, AOX was likely acquired from an oomycete-associated donor lineage along the stem leading to springtails, followed by repeated lineage-specific losses.

The respirometry experiments show that this horizontally acquired gene is physiologically active. In AOX-positive *F. candida* and *Proisotoma minuta*, mitochondrial oxygen consumption persisted after cytochrome-pathway inhibition and then declined after AOX inhibition. In contrast, AOX-negative species lost mitochondrial respiration after cytochrome-pathway inhibition alone, as expected for animals lacking an alternative terminal oxidase (43–45). This pharmacological signature strongly supports that springtail AOX has access to the mitochondrial ubiquinol pool and can operate as an alternative terminal oxidase when electron flow through complexes III and IV is constrained (25, 26, 29, 46, 47). Functional integration of this kind demonstrates that AOX is expressed, translated, targeted to mitochondria, imported into the inner membrane, folded into an active di-iron enzyme, and coordinated with host respiratory metabolism.

This functional result is important because AOX is best understood as a stress-tolerance respiratory pathway across eukaryotes. In plants, AOX contributes to metabolic and signaling homeostasis during abiotic and biotic stress and has been implicated specifically in low-oxygen and reoxygenation metabolism (21, 48). In fungi and protists, AOX can maintain respiration when the cytochrome pathway is limited and can contribute to stress responses, metabolic remodeling, and survival under bioenergetically challenging conditions (28–30, 49). In animals that naturally retain AOX, emerging work links the pathway to tolerance of sulfide, temperature, and hypoxia-reoxygenation stress (22, 23, 31, 50, 51). Thus, AOX acquisition in springtails appears to have restored a respiratory stress-tolerance function that had been lost in many animal lineages, including vertebrates and most arthropods.

An important question is why AOX would be advantageous under hypoxia or reoxygenation. The likely physiological advantage of springtail AOX is not increased ATP yield. AOX bypasses proton-translocating complexes III and IV and is therefore less energy conserving than the cytochrome pathway. Its advantage is more likely to be metabolic flexibility during respiratory stress (19, 21, 22, 48, 52–54). Here, I propose a potential mechanism of AOX-based metabolic flexibility where AOX functions in electron donor recycling needed for antioxidant metabolism and mitigation of excessive ROS production (Fig. 7). When oxygen is low or returns after hypoxia, cytochrome-pathway flux can become constrained, the ubiquinone pool and upstream electron donors can become highly reduced, and reactive oxygen species can increase (32, 55–58). AOX provides a route for oxidizing ubiquinol without engaging complexes III and IV, potentially protecting from electron-transfer-system over-reduction and helping maintain redox cycling for the tricarboxylic acid cycle and antioxidant metabolism.

**Figure 7.**
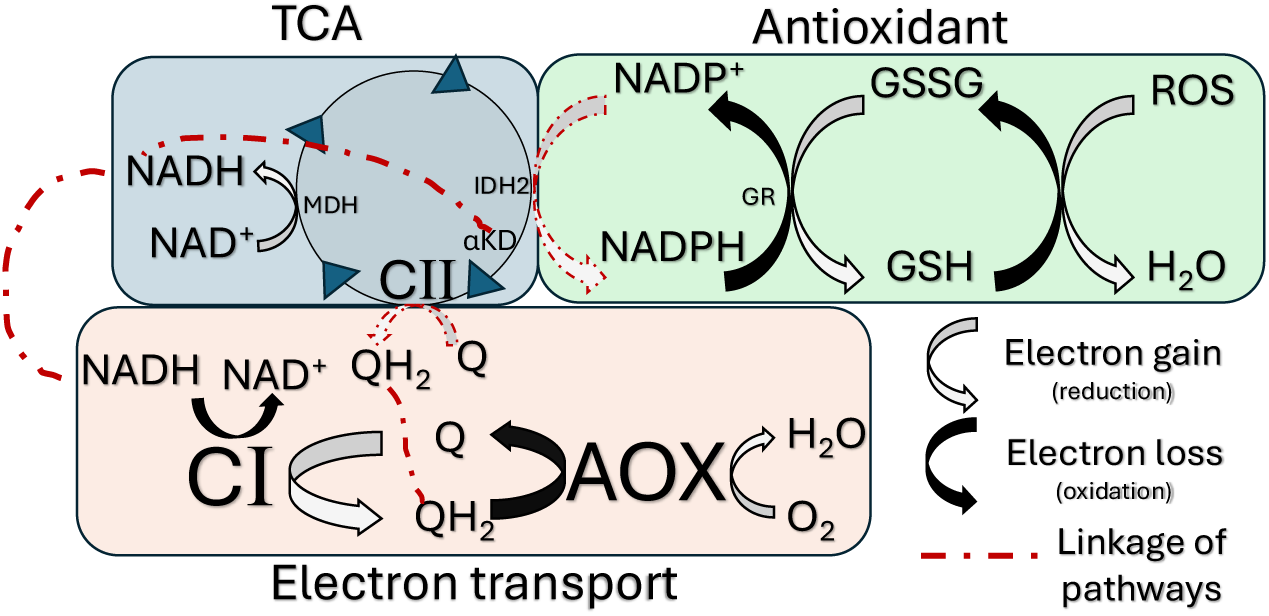
AOX-based respiration mediates the cellular redox state by driving electron donor recycling used by the TCA cycle and antioxidant metabolism. This non-ATP producing flux manages redox state when ATP demand is low or cytochrome pathway impairment. Malate dehydrogenase (MDH). Alpha-ketoglutarate dehydrogenase (αKD), isocitrate dehydrogenase 2 (IDH2), Glutathione reductase (GR), reduced (GSH) and oxidized (GSSG) glutathione.

The comparative results suggest that this metabolic function has been filtered by ecological conditions experienced across aboveground to belowground habitats. AOX was likely present near the origin of springtails but is now patchily distributed, with retention associated with belowground life forms and high hypoxia-exposure risk. This pattern is consistent with ancestral acquisition followed by repeated losses in lineages where AOX is weakly beneficial or unnecessary, and preferential retention where low or fluctuating oxygen makes AOX useful. The hypoxia-exposure experiments support this interpretation at the organismal level: AOX-positive species recovered robustly from hypoxia-reoxygenation, whereas AOX-negative species showed reduced or absent recovery. These experiments do not prove that AOX alone causes hypoxia tolerance, because species differ in many traits besides AOX. However, together with the respirometry and phylogenetic results, they support the hypothesis that AOX contributes to tolerance of hypoxia-reoxygenation stress.

The contrasting results for hemocyanin further suggest that oxygen-related traits can have different evolutionary dynamics. AOX presence tracked hypoxia-associated ecology more closely than hemocyanin presence, consistent with a direct role for AOX in modifying mitochondrial performance during cellular respiratory stress. Hemocyanin, by contrast, may reflect lineage history, expression regulation, oxygen affinity, concentration, or deployment rather than simple gene presence (42, 59). These differences suggest that hypoxia *tolerance* (via AOX) and hypoxia *avoidance* (a role for hemocyanin) may be supported by distinct trait classes with different evolutionary dynamics.

This study also strengthens the broader interpretation of AOX reacquisition in animals. Weaver and McDonald (18) identified putative lateral transfers of AOX into springtails, a fungus gnat, rotifers, and nematodes, with inferred fungal or protist donor lineages. Those cases suggested that AOX may be regained repeatedly in animals, but most remained sequence-based candidates requiring physiological validation. More recently Monesi et al. showed mitochondrial targeting and physiological integration (60) of horizontally acquired AOX in a fungus gnat (18).

Springtails now provide a genomic, functional, and ecological test case. The results of this study show that at least one putatively reacquired animal AOX has become a working mitochondrial pathway and is associated with environmental conditions under which AOX-like stress tolerance should be beneficial. This does not prove that all other putative animal AOX HGT events are functional or adaptive, but it raises the possibility that reacquisition of alternative oxidase respiration is a recurrent route to metabolic innovation in small animals exposed to harsh or fluctuating microenvironments. If so, this represents a remarkable example of parallel evolution of environmental respiratory stress tolerance acquired by evolutionarily distantly related animal lineages by multiple independent horizontal gene transfer events of a homologous mitochondrial gene from independent donors (18).

A key next step is to establish causality within AOX-positive springtails. The strongest tests would combine hypoxia-reoxygenation exposure with AOX-specific genetic or pharmacological perturbation, while measuring AOX protein abundance, mitochondrial ROS production, succinate dynamics, NADH/NAD+ balance, antioxidant capacity, and organismal recovery. Even with that limitation, the present results show that HGT can introduce metabolically important physiology into animals and that ecological context can determine where such physiology persists. In springtails, a foreign respiratory pathway appears to have become part of the bioenergetic basis of life belowground, linking gene acquisition to mitochondrial function, environmental tolerance, and soil biodiversity.

## Materials and Methods

Detailed experimental and computational procedures are provided in SI Appendix, SI Methods.

### Genome discovery and validation

Springtail genome assemblies (n = 202) and annotations were downloaded from NCBI. AOX candidates were detected with miniprot (61) using the *Folsomia candida* AOX protein as a reference and independently with tBLASTn. Candidate coding regions were reconstructed from aligned fragments, translated, clipped to the best stop-free AOX-like segment, and screened for the four conserved AOX iron-binding site motifs described in (18). Candidates recovering three or four motifs were treated as high confidence. For the *F. candida* RefSeq assembly (GCF_002217175.1), the AOX locus and a +/-200-kb genomic neighborhood were annotated and local protein-coding genes were classified by BLASTp against a reduced RefSeq database after removing *F. candida* self-hits. Microsynteny was tested by projecting the 16 protein-coding genes flanking AOX in the RefSeq assembly onto three additional *F. candida* assemblies with miniprot. Across AOX-positive genomes, AOX-centered windows were evaluated for contig position, sequence quality, rRNA annotations, and nearby gene-like intervals projected from a springtail protein reference panel.

### AOX phylogenetics

AOX protein sequences from springtails, oomycetes, fungi, plants, representative metazoans, and alphaproteobacteria were aligned and trimmed with ClipKIT (62). Maximum-likelihood phylogenies were inferred in IQ-TREE 3 (63) with ModelFinder, 1,000 ultrafast bootstrap replicates, and 1,000 SH-aLRT replicates. Donor-placement hypotheses were tested on a reduced curated alignment by comparing the unconstrained maximum-likelihood tree with topologies forcing springtail AOX as sister to either sampled oomycete AOXs or metazoan AOXs using approximately unbiased tests with 10,000 RELL replicates.

### Respirometry and hypoxia assays

*F. candida, Proisotoma minuta, Pseudosinella violenta,* and *Isotoma delta* were maintained in laboratory cultures. Mitochondrial preparations were obtained from each species and assayed at 25 degrees C with an Oroboros O2k high-resolution respirometer. Respiration was stimulated with pyruvate, malate, ADP, glutamate, succinate, cytochrome c, and CCCP. Antimycin A was used to inhibit the cytochrome pathway, salicylhydroxamic acid to inhibit AOX, and rotenone to estimate residual oxygen consumption for background correction. Oxygen fluxes were normalized to protein content and expressed as absolute rates and flux control ratios. Hypoxia tolerance was assayed by exposing individuals to 1% O2 at 99% relative humidity and 23 degrees C for 4 h or 16 h, followed by reoxygenation. Recovery was scored as normal activity versus no movement 24 h after reoxygenation, with normoxic chamber controls. Species and treatment effects were tested with Fisher’s exact tests and Firth penalized logistic regression.

### Comparative analyses

AOX presence, habitat association, hypoxia-exposure category, and hemocyanin presence were scored for species represented in a published genome-scale springtail phylogeny (Table S1). Habitat categories followed published springtail life-form classifications, and hypoxia exposure was assigned from life form and reported microhabitat occupancy (38, 39, 64). Hemocyanin homologs were identified with tBLASTn, HMM profiles, phylogenetic analyses, and structural comparisons for ambiguous type-3 copper protein candidates. AOX and hemocyanin ancestral states were reconstructed with discrete-state Mk models and stochastic character mapping in phytools (65). Correlated evolution was tested with Pagel’s discrete models (66), and ecological predictors of AOX or hemocyanin presence were evaluated with phylogenetic logistic regression in phylolm (66).

## Acknowledgments

I am grateful for thoughtful feedback from Jonathan Wendel, Dennis Lavrov, Nicole Valenzuela, and Dean Adams on earlier drafts of this manuscript. Springtail culturing was supported by Brooklyn Damgaard; Blake Stanton assisted with literature review. This project was supported by faculty startup funds provided by the Department of Ecology, Evolution, and Organismal Biology, the College of Agriculture and Life Sciences at Iowa State University.

## Author Contributions

R.J.W. designed research, performed research, analyzed data, and wrote the paper.

## Supporting Information

### SI Methods

#### AOX genomic context in the *Folsomia candida* RefSeq genome

The *F. candida* AOX validation workflow used the publicly available annotated RefSeq assembly GCF_002217175.1 / ASM221717v1. The original AOX discovery interval was treated as NW_019091241.1:119,206-120,320, whereas gene-structure, targeting, and local-context analyses used the complete overlapping RefSeq gene model LOC110862114. This model is annotated as alternative oxidase, mitochondrial-like, with transcript XM_022111270.2 and protein XP_021966962.1, and spans NW_019091241.1:119,078-121,029 on the minus strand. A partially overlapping opposite-strand lncRNA annotation, LOC118433373, was recorded separately so that the AOX protein-coding model was not conflated with a noncoding feature.

Genome FASTA, GFF3, protein FASTA, CDS, RNA, and GenBank-format records were obtained from the NCBI/RefSeq assembly package. Exon, CDS, mRNA, and gene features for LOC110862114 were parsed from the genomic GFF3 by filtering for the AOX gene and transcript identifiers. Intron intervals were inferred from gaps between adjacent exon coordinates after sorting exons in genomic order. Because AOX is encoded on the minus strand, splice donor and acceptor motifs were extracted from the genomic FASTA and reverse-complemented into transcript orientation before classifying splice boundaries.

To characterize local genomic context, a nominal +/-200 kb neighborhood centered on the AOX gene was extracted from the RefSeq assembly. Because AOX lies near the beginning of NW_019091241.1, the window was clipped at the scaffold start, yielding NW_019091241.1:1-321,029. Gene density was summarized by counting annotated gene features within this clipped window. Protein identifiers overlapping the window were extracted from CDS features and used to retrieve protein sequences for local taxonomic-coherence analyses.

Neighborhood proteins were compared to NCBI protein databases using the bundled BLAST workflow. Hits assigned to *F. candida* were removed for the best non-self summaries, and BLAST hit taxids were resolved with NCBI Taxonomy. The retained hit tables were grouped as Metazoa, non-metazoan eukaryote, prokaryote, or unknown. The central contamination test was whether non-metazoan eukaryote affinity was distributed broadly across the local scaffold or restricted to the AOX query itself. Eukaryotic rRNA features were predicted across the assembly with barrnap --kingdom euk, and the resulting GFF was searched for features on NW_019091241.1 and within the AOX-centered neighborhood.

Public paired-end WGS reads from SRA run SRR2743547 were used as an independent assembly-support check. Reads were mapped to the RefSeq assembly with bwa-mem2, sorted and indexed with samtools (1), and summarized with samtools flagstat and samtools stats. Genome-wide and scaffold-level 1-kb coverage were computed with mosdepth; per-base depth across the AOX discovery interval was extracted with samtools depth. Read-pair behavior was evaluated globally with flagstat, locally with a region-specific proper-pair and mate-reference summary, and globally for insert-size distribution with Picard CollectInsertSizeMetrics.

#### AOX iron-binding-site mapping and mitochondrial targeting

The AOX protein sequence XP_021966962.1 was extracted from the RefSeq protein FASTA. Four conserved AOX iron-binding-site motifs were mapped from amino-acid coordinates to genomic coordinates by concatenating CDS segments in transcript order while accounting for minus-strand orientation and GFF phase. Because exons include untranslated sequence but the motifs occur in the coding region, motif coordinates were mapped to CDS spans rather than to whole-exon spans. For gene-model visualization, UTR-only portions of exons and CDS portions of mixed exons were treated as separate graphical features.

Mitochondrial-targeting analyses used a comparative AOX set containing *Pythium aphanidermatum* AOX (CAE11918.1), *F. candida* AOX (XP_021966962.1), *Crassostrea virginica* AOX-like protein (XP_022314634.1), and *Branchiostoma floridae* AOX-like protein (XP_035700253.1). Four FASTA inputs were prepared: full-length proteins, N70 proteins containing the first 70 amino acids, N120 proteins containing the first 120 amino acids, and deltaN50 proteins with the first 50 amino acids removed. TargetP 2.0 (2) was run in non-plant mode, recording predicted class, prediction score, cleavage-site position, local cleavage motif, and cleavage-site probability when provided. The deltaN50 construct tested whether TargetP identified an internal-like signal after removal of the N terminus. ATP5B and HSP60 comparison sets were run as mitochondrial positive controls

#### AOX microsynteny among *F. candida* genomes

The AOX microsynteny workflow was designed as an AOX-centered local genomic-context analysis, not chromosome-scale synteny. The RefSeq assembly was used as the reference genomic context. AOX was anchored by protein XP_021966962.1 / gene LOC110862114, and the local RefSeq neighborhood was defined as AOX plus the eight nearest protein-coding genes on each coordinate side of AOX. This produced a 17-marker reference set ranked from -8 to +8 relative to AOX. Because AOX is on the minus strand, the workflow side labels are coordinate-based left/right flanks rather than promoter-direction upstream/downstream labels.

Three additional *F. candida* assemblies with resolvable genome FASTA files were analyzed. AOX anchors in the target assemblies were obtained from the unified AOX scan outputs using the configured anchor-priority scheme. In the main run, all three targets were anchored from tblastn fragment coordinates. For each target, the workflow extracted a 400,001 bp AOX-centered window around the AOX anchor midpoint and projected the 17 RefSeq marker proteins onto the target window with miniprot (3). The best hit per query marker was retained, converted back to absolute target coordinates, assigned its expected side from the RefSeq marker table, and scored relative to the target AOX midpoint.

Support tiers were assigned from AOX anchoring and non-AOX flank recovery. Tier1_strong required a recovered AOX anchor, at least three projected non-AOX flanks, and at least two projected non-AOX flanks on the expected side of AOX. Sensitivity analyses repeated the projection with alternative target-region sizes (100 kb, 200 kb, 500 kb, and whole contig) and alternative anchor choices. Fragment and model anchors provided independent support; stitched-anchor configurations reproduced the final result but fell back to fragment anchors in the collated summaries, so they are treated as fallback confirmations rather than independent stitched-coordinate validations. A same-contig negative-control window far from AOX and an independent tblastn sanity check were used as additional QA checks.

#### AOX scan of 202 springtail genome assemblies and flanking-interval workflow

The accession-wide AOX workflow scanned 202 springtail genome assemblies. NCBI genome datasets were downloaded and unpacked into a standardized run-root structure driven by an accession manifest and project configuration file. Stage 1 aligned the *F. candida* AOX protein XP_021966962.1 to each assembly with miniprot. Pairwise mapping format (PAF) output was parsed to record query coverage, percent identity, aligned amino-acid span, mapping quality, contig, strand, and genomic coordinates. A candidate was provisionally called AOX-positive when it met all configured Stage 1 thresholds: query fraction >= 0.30, identity >= 30%, and aligned AOX span >= 80 amino acids. Candidate ranking favored higher query fraction, identity, aligned span, and mapping quality.

For each AOX-positive accession, the best AOX interval was expanded to a +/-200 kb window, clipped to contig boundaries. Sequence-context variables included contig length, window length, distance from AOX to the nearest contig edge, N fraction, softmasked fraction, and rRNA predictions from barrnap. The locus-context score was a coarse triage screen rather than a definitive contamination test. Scores increased with stronger AOX alignment, higher identity, low N content, modest softmask fraction, absence of rRNA in the AOX window, greater rRNA distance, and greater distance from contig edges. Scores decreased for short AOX spans, high N content, high repeat/softmask fraction, rRNA in or near the window, and edge-proximal loci. Final context classes were clean_likely_authentic, needs_manual_inspection, and suspicious_or_low_confidence.

Because many assemblies were fragmented, the workflow added a protein-centric recovery layer. miniprot was rerun in GFF mode with j1 and j2 settings, CDS features were extracted and translated, and translated models were clipped to the best stop-free AOX-like segment. AOX-positive assemblies were also searched independently with tblastn using the *F. candida* AOX protein. The configured tblastn filters were minimum bitscore 50, maximum E value 1e-5, minimum amino-acid fragment length 25, and up to 100 targets. High-scoring segment pairs were translated in subject orientation, reduced to stop-free peptide fragments, deduplicated when strongly overlapping, and stitched into candidate reconstructions only when fragments occurred on the same contig and strand with compatible AOX-reference and genomic order. Overlaps in reference space were trimmed; gaps were represented with X characters.

Stitched candidates were screened for four AOX iron-binding-site motifs defined from the *F. candida* AOX reference: IBS1 = LE[TS], IBS2 = NERMHL, IBS3 = GYLEEE, and IBS4 = RADE..H (4). Candidates recovering three or four motifs were classified as high_confidence; candidates recovering zero to two motifs were retained as low_confidence sequence evidence. The authoritative high-confidence count in the final packet is 65 high-confidence stitched candidates among 75 AOX-positive accessions.

I also tested whether AOX-positive loci were embedded in broader springtail-like sequence context. A reference protein panel was built from annotated *F. candida* and *Allacma fusca* proteomes. For each AOX-positive accession, the +/-200 kb AOX window from Stage 1 was used as a local target, and miniprot projected all reference proteins into that window. Projections were retained as pass_local when query fraction was >= 0.30, identity was >= 35%, and aligned reference span was >= 80 amino acids. Each projection was classified as AOX_overlap if it intersected the AOX interval, or as left/right of AOX using midpoint offset from the AOX interval. Redundant pass_local projections were collapsed first into family-level local loci and then into distinct interval-level loci by merging overlapping or near-adjacent family loci. Final per-accession summaries recorded distinct left, right, and AOX-overlap intervals and nearest left/right distances.

#### AOX phylogeny and topology testing

The broad AOX context phylogeny used a 165-aa aligned AOX protein file containing 172 sequences before trimming, including 26 springtail AOXs, 6 oomycetes, other metazoans, fungi, plants/green algae, alphaproteobacteria, and other non-metazoan eukaryotic representatives (4). Springtail AOX sequences were drawn from the high-confidence springtail AOX set derived from tblastn-stitched candidates. The alignment was trimmed with ClipKIT (5) in smart-gap mode, yielding 172 sequences and 164 retained amino-acid sites. IQ-TREE 3.1.1 (6) was run with ModelFinder (7), 10 independent tree-search runs, 1,000 ultrafast bootstrap replicates, and 1,000 SH-aLRT replicates. ModelFinder selected LG+I+R7 by BIC for the primary broad context tree.

Because the broad tree was taxon-rich relative to the short retained alignment, it was treated as a context tree rather than the formal donor-placement test. A C20 mixture-model guide-tree sensitivity analysis was run under LG+C20+F+G. The guide tree completed and recovered the same qualitative focal relationship.

Formal topology tests used a reduced manually curated donor-test alignment containing 36 AOX proteins before trimming: 12 springtails, 6 oomycetes, 12 non-panarthropod metazoans, and 6 alphaproteobacteria used to anchor the comparisons. The reduced alignment was trimmed with ClipKIT smart-gap mode and analyzed in IQ-TREE with ModelFinder, 10 independent tree-search runs, 1,000 ultrafast bootstrap replicates, and 1,000 SH-aLRT replicates (8). The best-fit model for the reduced tree was LG+I+G4 selected by BIC.

Approximately unbiased (AU) topology tests were then run in IQ-TREE using 10,000 RELL resamplings. Three candidate trees were compared in a verified line order: the unconstrained reduced maximum-likelihood tree, a topology constraining springtail AOXs as sister to sampled oomycete AOXs, and a topology constraining springtail AOXs as sister to sampled non-panarthropod metazoan AOXs. Constraint trees were anchored so that the springtail clade, tested sister group, competing group, and alphaproteobacterial anchors were represented explicitly. A verification script checked that candidate_trees.nwk line order matched the intended tree identities before interpretation.

#### Springtail cultures

Laboratory cultures of *F. candida*, *Proisotoma minuta*, and *Pseudosinella violenta* were established from individuals purchased from a commercial supplier (Springtails US, IL). *Isotoma delta* cultures were started from individuals collected around Bessey Hall on the Iowa State University campus. Cultures were maintained at approximately 23 degrees C in small plastic containers filled with natural charcoal (*F. candida*, *P. minuta*, and *P. violenta*) or potting soil mix (*I. delta*; Burpee organic). Cultures were supplied with dried nutritional yeast (Bragg, CA) twice weekly. Humidity in charcoal-based cultures was maintained by adding distilled water to form approximately a 1 cm pool at the bottom of the container. Soil cultures had mesh-covered ventilation holes and were misted with distilled water twice weekly. Relative humidity in culture containers was not measured.

#### Electron-transfer-system substrates, uncoupler, and inhibitors

The respirometry protocol used pyruvate (P; 5 mM), malate (M; 2 mM), glutamate (G; 10 mM), succinate (S; 10 mM), ADP (D; 2.5 mM), and cytochrome c (c; 10 uM) to stimulate respiratory states. CCCP (U; 0.5 uM) was used as the uncoupler. Antimycin A (AA; 2.5 uM) was used to inhibit the cytochrome pathway, salicylhydroxamic acid (SHAM; 1 mM) to inhibit AOX, and rotenone (0.5 uM) to inhibit complex I and estimate residual oxygen consumption. Substrates were dissolved in deionized water; uncoupler and inhibitors were dissolved in 100% ethanol.

#### High-resolution O2k respirometry and statistical analysis

Mitochondrial preparations were obtained from samples of approximately 200 springtails homogenized on ice in 1 mL mitochondrial isolation buffer (250 mM sucrose, 5 mM Tris-HCl, 2 mM EGTA, 1% w/v fatty acid-free BSA, pH 7.4) using a glass Potter-Elvehjem tissue grinder. Samples were homogenized by 5-6 manual strokes with rotation of a tight-fitting PTFE pestle. Homogenates were passed through Miracloth to remove cellular debris and kept on ice before respirometry. Protein content was measured with a Qubit 3 fluorometer (Invitrogen), and remaining homogenate was archived at -80 degrees C.

Oxygen consumption was measured with an Oroboros O2k high-resolution respirometer (Oroboros Instruments, Innsbruck, Austria) in 2 mL chambers containing MiR05 respiration medium (110 mM sucrose, 60 mM lactobionic acid, 20 mM taurine, 20 mM HEPES, 10 mM KH2PO4, 3 mM MgCl2, 0.5 mM EGTA, 1% BSA, pH 7.1). The processed O2k workbook recorded a modified SUIT-008 O2 mt D026 protocol, 2 mL chamber volume for all sample blocks, and run temperatures of 23 or 25 degrees C. After addition of mitochondrial preparation to the chamber, substrates, uncoupler, and inhibitors were titrated sequentially. Pyruvate and malate stimulated the LEAK state through complex I, ADP initiated OXPHOS, cytochrome c tested for preparation-related electron-carrier limitation, glutamate supported the PMG pathway, succinate engaged complex II, and uncoupler titration estimated maximal respiration.

AOX functionality was tested pharmacologically. In the main control sequence, antimycin A was titrated to inhibit the cytochrome pathway. Persistence of oxygen flux after antimycin A was interpreted as cytochrome-pathway-inhibitor-resistant respiration, and subsequent SHAM titration tested whether this remaining flux was AOX sensitive.

Reciprocal pretreatment assays titrated SHAM or antimycin A at the beginning of the protocol after mitochondrial preparation was added to the chambers. Oxygen consumption values were extracted from DatLab/O2k outputs and curated into the final analysis table. Flux control ratios were calculated by dividing each retained respiratory state by the uncoupled maximum-respiration state so that maximum respiration was scaled to 1.0.

The final R workflow reshaped mass-specific flux variables into long format, retained the primary respiratory states used for plotting and statistics, calculated flux control ratios, and generated OXPHOS-capacity contrasts. For each retained respiratory state within the relevant treatment subset, one-way ANOVA tested species effects and Tukey HSD was used for post hoc comparisons and compact-letter displays. The workflow reported mean, standard error, HSD letters, ANOVA F statistics, degrees of freedom, and P values. Figure error bars were generated from the plotting code and correspond to the statistic indicated in each figure legend or table.

#### Hypoxia exposure assay

Hypoxia tolerance was assayed by exposing springtails to 1% O2, 99% relative humidity, and 23 degrees C in a sealed chamber for 4 h or 16 h. Individuals were placed in wells of a 48-well plate and reoxygenated after the exposure period. Activity was assessed at 0.5, 2, 4, and 24 h after reoxygenation, with the primary endpoint scored at 24 h as recovered (normal activity) versus not recovered (no movement). Normoxic chamber controls were run under the same handling conditions to evaluate chamber and handling effects. Species and treatment effects were tested with Fisher exact tests and Firth penalized logistic regression because recovery outcomes showed complete or near-complete separation among species and treatments.

#### Springtail trait data for comparative phylogenetics

Comparative analyses used the 47-tip ASTRAL species tree from the Yu et al. (9) genome-scale springtail phylogeny as a fixed phylogenetic backbone. Species names in the tree were matched to the trait table by the species field.

AOX analyses excluded the two non-Collembola outgroups and three taxa without AOX state calls, yielding a 42-taxon Collembola AOX analysis set. AOX presence or absence was determined using tblastn searches with a fungal AOX protein sequence (XP_006680383.1) as the query, following (4). AOX presence was converted to a binary trait with present coded as 1 and absent coded as 0.

Habitat association was classified as epedaphic, hemiedaphic, or euedaphic based on Potapov et al. (10), Li et al. (11), and the database for soil invertebrate biological and ecological traits (12). Hypoxia exposure risk was assigned based on each species’ life form and reported microhabitat occupancy. For example, euedaphic species were considered likely to experience recurrent hypoxia because oxygen availability generally declines with soil depth, although by how much depends on soil structure, moisture, and respiration (13–15). However, hypoxia exposure risk was not assumed to map perfectly onto life form alone. Some above ground-associated microhabitats, such as the interiors of decomposing logs may also experience low oxygen conditions, whereas some below ground habitats, such as ant nests with surface connections, may remain sufficiently ventilated to avoid substantial hypoxia (16, 17). Hypoxia exposure was scored as recurrent exposure and avoidance potential rather than as direct oxygen concentration. Low was assigned to surface or readily escapable microhabitats, Moderate to lower-litter, moss, decaying wood, under-stone, ant-nest, cave-passage, or upper-soil interfaces, and High to mineral soil, deeper humus/soil pore spaces, saturated or interstitial substrates, and eudaphic morphologies with limited escape.

The sensitivity workflow repeated the comparative analyses with avoidance-aware alternative habitat and hypoxia calls from alt.habitat and alt.hypoxia. These alternative calls reduced the tendency to score behaviorally escapable belowground or interface habitats as High hypoxia. The sensitivity rerun used the same tree, AOX states, and hemocyanin calls as the primary workflow, changing only the ecological predictor columns.

#### Hemocyanin identification and structure-informed adjudication

Hemocyanin homologs were identified with a genome-based and structure-assisted workflow that explicitly accounted for the close relationship between hemocyanins and phenoloxidases, both arthropod type-3 copper proteins. Public genome assemblies were downloaded and converted to genome FASTA files. Hemocyanin and phenoloxidase reference FASTAs were used as competing tblastn query sets against local nucleotide BLAST databases for each assembly. First-pass summaries recorded the best hemocyanin-like and phenoloxidase-like hits, bit scores, E values, alignment lengths, percent identity, and bit-score differences. Strong hemocyanin-like sequence similarity was not treated as definitive hemocyanin evidence when phenoloxidase-like evidence was stronger.

Candidate loci were extracted from genome assemblies using tblastn hit coordinates and flanking windows. Open reading frames from these candidate loci were scored against HMM profiles (18) for broad hemocyanin, phenoloxidase, springtail hemocyanin subtype Hc1, springtail hemocyanin subtype Hc2, and PPO. A small protein tree containing selected candidate proteins and reference sequences was used as an additional adjudication layer to identify Hc-like, PPO-like, and ambiguous placements. For ambiguous taxa, full-ish Hc/PPO reference proteins were aligned back to genome FASTAs with miniprot; CDS features were translated directly from miniprot GFF output and rescored against the HMM profiles.

A curated structure panel was predicted with ColabFold 1.5.5 (19) using an Apptainer/SIF workflow on NOVA HPC cluster at Iowa State Unviersity. Candidate and reference models were compared with Foldseek (20). For each candidate model, the workflow summarized the best Hc structural score, best PPO structural score, Hc-minus-PPO structural delta, top reference class, top reference-class counts, and model confidence. Final species-level hemocyanin calls merged sequence/HMM evidence, miniprot rescue, protein-tree placement, model confidence, Foldseek structural neighborhood, and prior call state. Structure alone was not used to call absence, and mixed Hc/PPO structural neighborhoods were treated conservatively. Candidate_present species with strong Hc structural and tree support were upgraded to present; uncertain species with strong or moderate Hc structural evidence and Hc-favoring HMM scores were upgraded to candidate_present. Conservative binary coding treated present plus candidate_present as hemocyanin-positive when used in analyses. Uncertain and no-data states were excluded or handled as ambiguous for focal binary tests.

#### Ancestral-state reconstruction, correlated evolution, and phylogenetic logistic regression

AOX ancestral states were reconstructed on the rooted species phylogeny using discrete-state continuous-time Markov models implemented in phytools (21). Equal-rates and all-rates-different transition models were fit with alternative root priors. For AOX, root priors included flat (equal likelihood presence, absence), absence-biased 0.05 present, 0.95 absent), and fixed-absence priors (0 present, 1 absent), with the fixed AOX-absence root prior treated as biologically relevant because AOX is absent from the sampled non-Collembola outgroups and from most arthropods (4). Hemocyanin ancestral-state reconstruction used analogous binary-state model fitting; taxa no or low confidence hemocyanin data were excluded from the analysis.

Pagel independent-versus-dependent models were fit with phytools::fitPagel. AOX was tested against high hypoxia, eudaphic habitat, and inclusive hemocyanin presence. Hemocyanin was tested against moderate/high hypoxia and AOX. Model support was evaluated with likelihood-ratio tests and AIC differences. The sensitivity workflow exported parallel Pagel summaries and Q matrices using the alternative ecological predictor columns; the matching sensitivity AOX-vs-hypoxia and AOX-vs-habitat Pagel results are expected because the High hypoxia and EUD habitat binary vectors are identical after revised scoring.

Phylogenetic logistic regressions were fit with phylolm::phyloglm using method = logistic_MPLE (22). Primary and sensitivity workflows modeled AOX presence as a function of hypoxia category and habitat category. Hemocyanin was analyzed in parallel as the response trait and hypoxia exposure and habitat as predictors. Coefficient tables report log-odds estimates, standard errors, z statistics, P values, approximate confidence intervals, and exponentiated odds ratios.

### Supplementary Results

The *F. candida* AOX locus is supported by several independent lines of genomic evidence. The AOX discovery interval is contained within RefSeq gene LOC110862114, which has a multi-exon nuclear gene model, canonical GT-AG splice boundaries in transcript orientation, and RefSeq RNA-seq model evidence supporting all annotated introns (Fig. S1A). The AOX-containing scaffold has near-genome-median WGS depth, continuous nonzero per-base coverage across the AOX discovery interval, and normal local read-pair behavior (Fig. S1C). The local neighborhood is overwhelmingly host-like: non-metazoan eukaryote BLAST/taxonomy rows are restricted to AOX-associated hits, and barrnap detected no rRNA feature on the AOX-containing scaffold (Fig. S1B). TargetP 2.0 predicts an N-terminal mitochondrial targeting signal for *F. candida* AOX, and the deltaN50 truncation removes this TargetP signal (Fig. S1C).

AOX microsynteny across *F. candida* assemblies further supports an endogenous locus. All three target *F. candida* assemblies recovered AOX and all 16 non-AOX RefSeq flanking markers on the expected side of AOX, with Tier1_strong support in each assembly (Fig. S2A,B). The result remained stable across target-window and anchor-sensitivity analyses, was reproduced by fragment-based and model-based AOX anchoring, was supported by an independent tblastn sanity check, and was absent from a same-contig negative-control window.

Across the 202-accession springtail genome scan, Stage 1 identified 75 AOX-positive accessions. Context screening classified 20 as clean_likely_authentic, 45 as needs_manual_inspection, and 10 as suspicious_or_low_confidence; these are triage classes rather than automatic acceptance/rejection labels. No AOX-positive window contained an rRNA feature. The tblastn/stitched-candidate layer recovered 65 high-confidence stitched AOX candidates among the 75 positives, and Stage 2B local-context projection recovered 608 deduplicated local intervals from 6,898 pass_local projections. Because many assemblies are fragmented and edge-proximal, the Stage 2B interval results are interpreted as fragment-aware local-context support rather than strict macrosynteny.

AOX phylogenetic analyses placed high-confidence springtail AOX sequences in an oomycete-associated region of AOX tree space. The broad 172-sequence context tree recovered a springtail clade and a springtail-plus-oomycete focal grouping, and the completed LG+C20+F+G guide tree recovered the same qualitative focal relationship. Formal reduced AU tests retained the oomycete-sister topology (deltaL = 0.0942; p-AU = 0.539) and rejected the non-panarthropod metazoan-sister topology (deltaL = 18.186; p-AU = 0.00805). These tests support an oomycete/protist-associated origin of springtail AOX relative to the sampled alternatives, but they do not identify a single exact donor lineage.

O2k respirometry supports functional integration of AOX in AOX-positive springtails. Under control conditions, *F. candida* and *P. minuta* showed AOX-associated oxygen flux whereas AOX-negative *P. violenta* and *I. delta* did not. AOX-associated flux represented approximately one-sixth of maximal respiration in the AOX-positive species.

Antimycin A treatment supported cytochrome-pathway-inhibitor-resistant respiration in AOX-positive species, and SHAM treatment removed the AOX-specific respiratory signal while leaving cytochrome-pathway respiration available. Hypoxia-reoxygenation assays showed high recovery in AOX-positive species and reduced or absent recovery in AOX-negative species under the tested conditions.

Comparative phylogenetic analyses support preferential AOX retention in hypoxia-prone and belowground lineages. Primary phylogenetic logistic regressions and Pagel tests supported positive associations between AOX presence and High hypoxia/EUD habitat, and the avoidance-aware sensitivity rerun preserved the same qualitative conclusions while reducing some effect sizes. Hemocyanin gene-presence analyses did not show the same ecological pattern, and structure-informed hemocyanin adjudication reduced uncertainty without converting hemocyanin presence into a significant predictor of AOX or hypoxia-associated ecology.

## Figures

**Figure S1.**
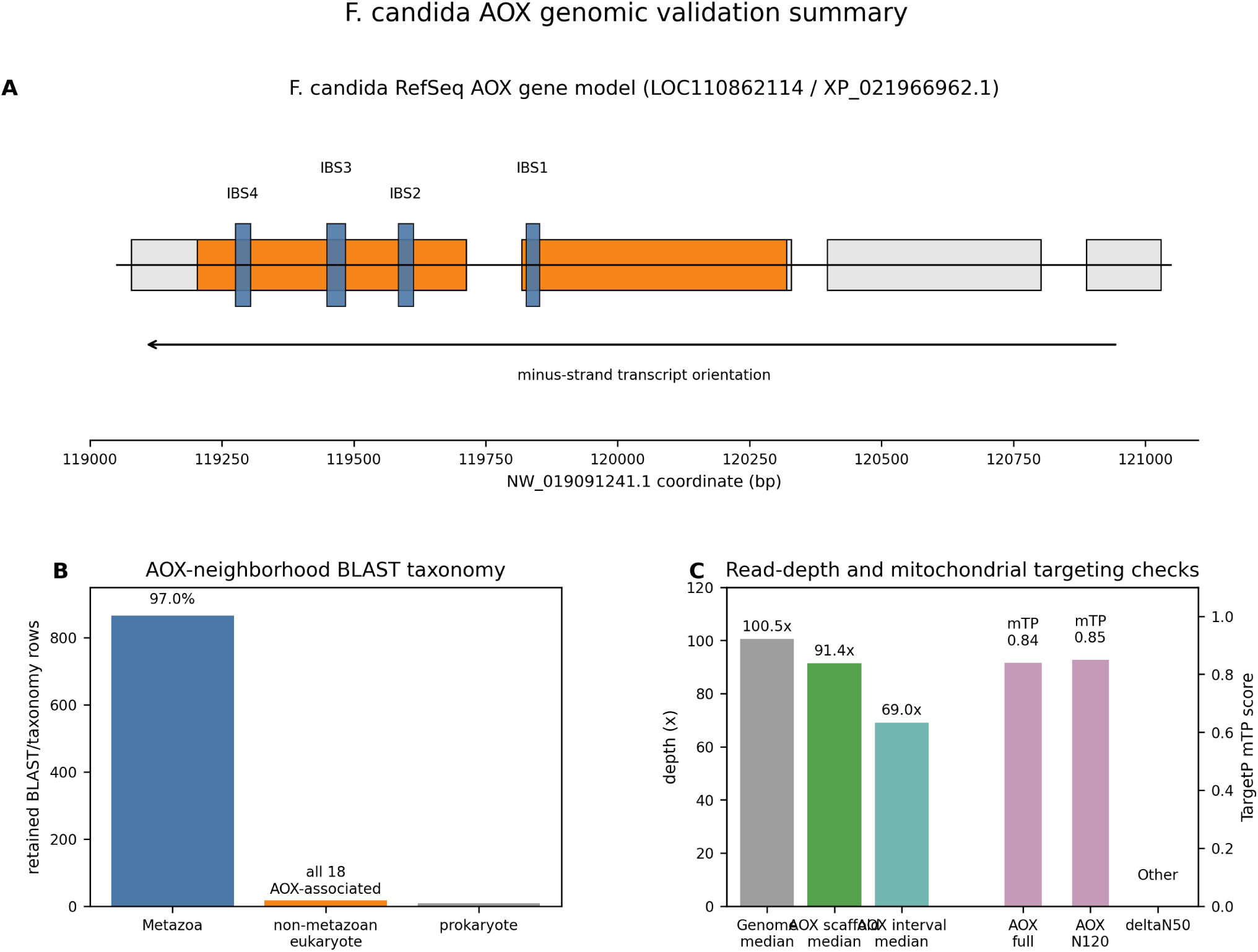
Genomic validation summary for *F. candida* AOX. (A) RefSeq AOX gene model showing UTR and CDS portions of exons and the four mapped iron-binding-site motifs. The transcript is on the minus strand. (B) Taxonomic summary of retained BLAST/taxonomy rows for proteins in the AOX neighborhood. Non-metazoan eukaryote rows were restricted to AOX-associated hits. (C) WGS read-depth and TargetP checks. AOX scaffold depth is near the genome median, and TargetP predicts a mitochondrial targeting peptide in full-length and N120 AOX but not after removal of the first 50 amino acids.

**Figure S2.**
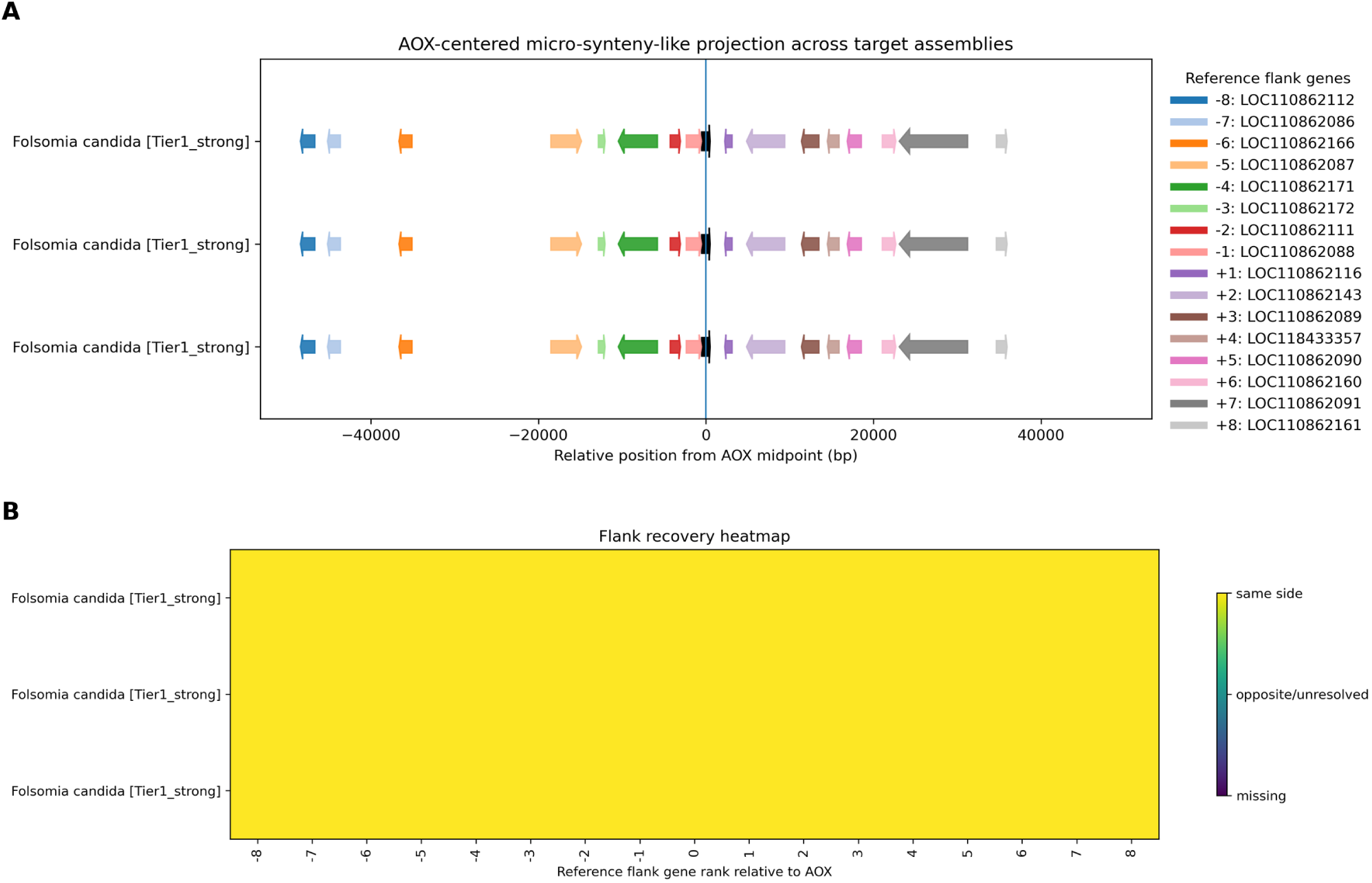
*Folsomia candida* AOX microsynteny validation. (A) AOX-centered projection of the 16 non-AOX RefSeq flanking markers onto three additional *F. candida* assemblies. AOX is centered at position 0, and each projected marker is shown by relative position from the AOX midpoint. All three targets were classified as Tier1_strong. (B) Flank-recovery heatmap showing same-side recovery for every ranked reference marker from -8 to +8 in each target assembly.

**Figure S3.**
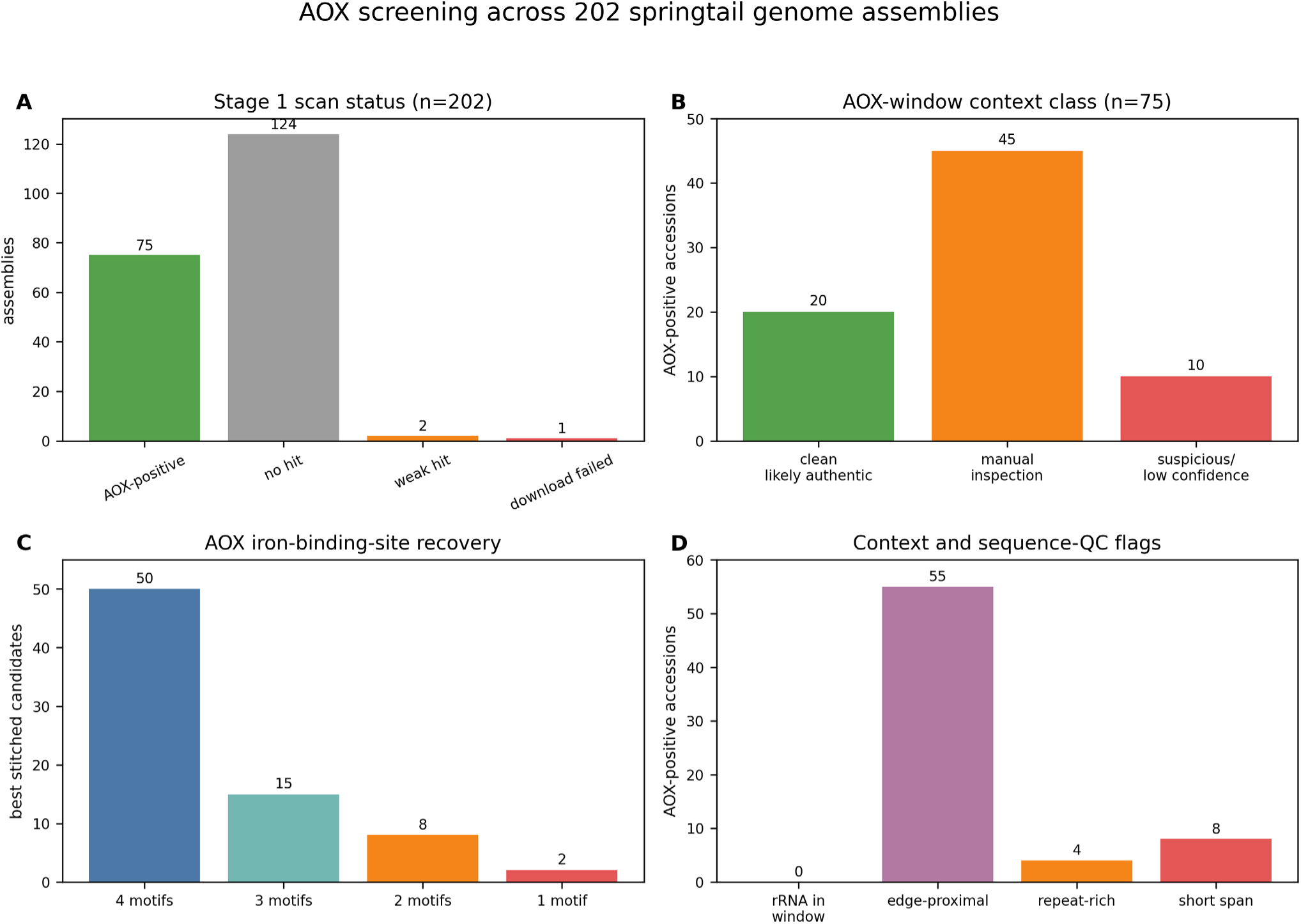
AOX screening across 202 springtail genome assemblies. Panels summarize Stage 1 scan status, context-class triage among AOX-positive accessions, diagnostic iron-binding-site recovery in best stitched candidates, and key sequence/context flags.

**Figure S4.**
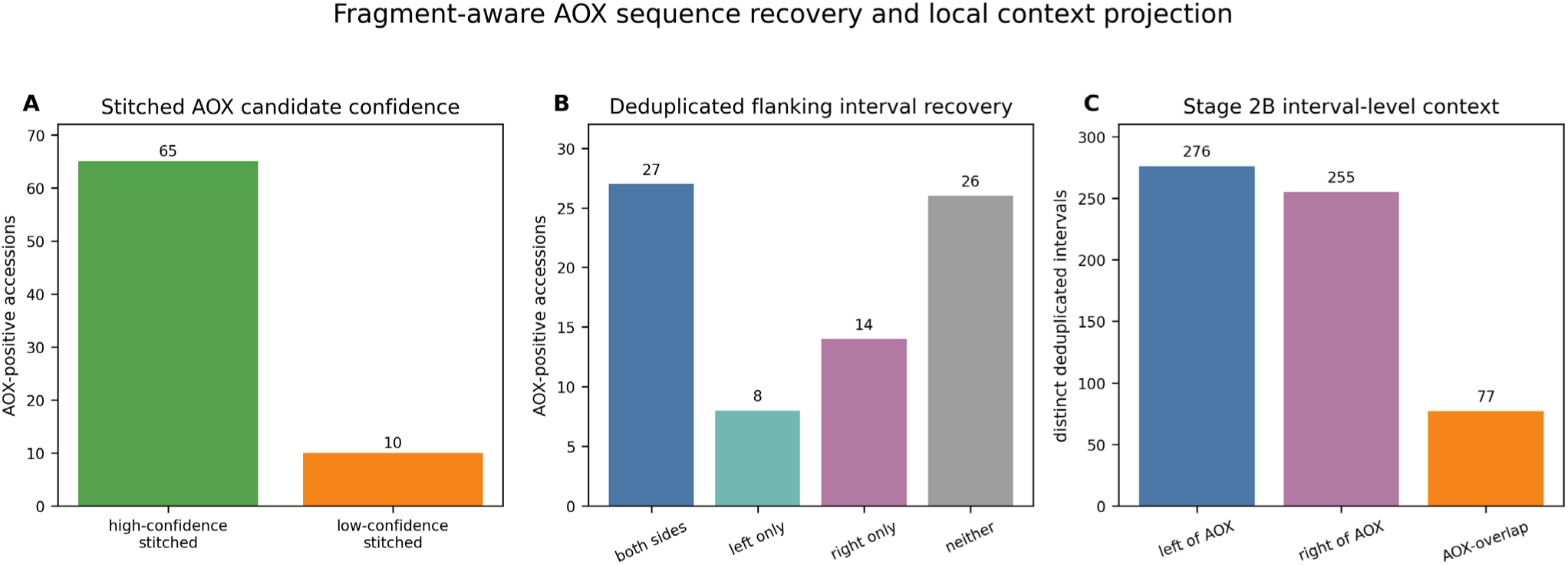
Fragment-aware AOX sequence recovery and local context projection. (A) High-confidence and low-confidence stitched AOX candidate counts among AOX-positive accessions. (B) Recovery of deduplicated flanking intervals on both sides, one side, or neither side of AOX. (C) Stage 2B distinct interval counts by side relative to AOX.

**Figure S5.**
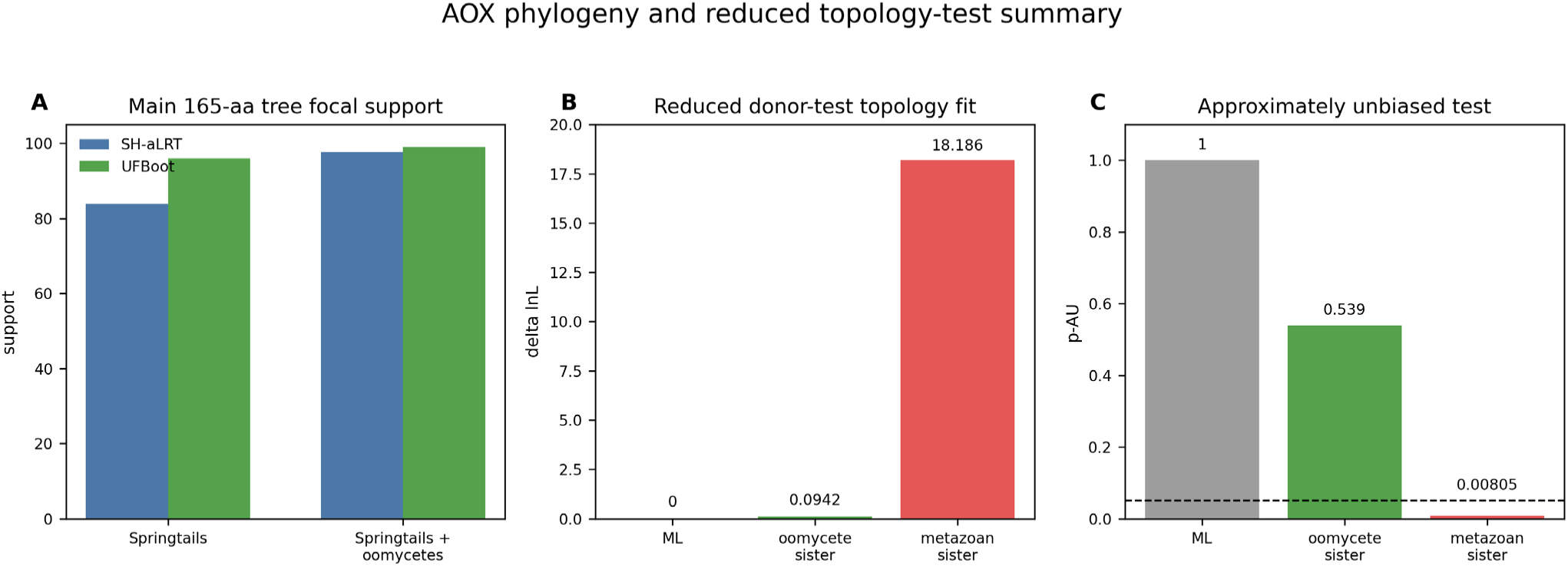
AOX phylogeny and reduced topology-test summary. (A) Focal support values in the broad 165-aa AOX context tree. (B) Delta log-likelihood for reduced donor-test candidate topologies. (C) Approximately unbiased test p-values. The metazoan-sister topology was rejected, whereas the unconstrained and oomycete-sister topologies were retained.

**Figure S6.**
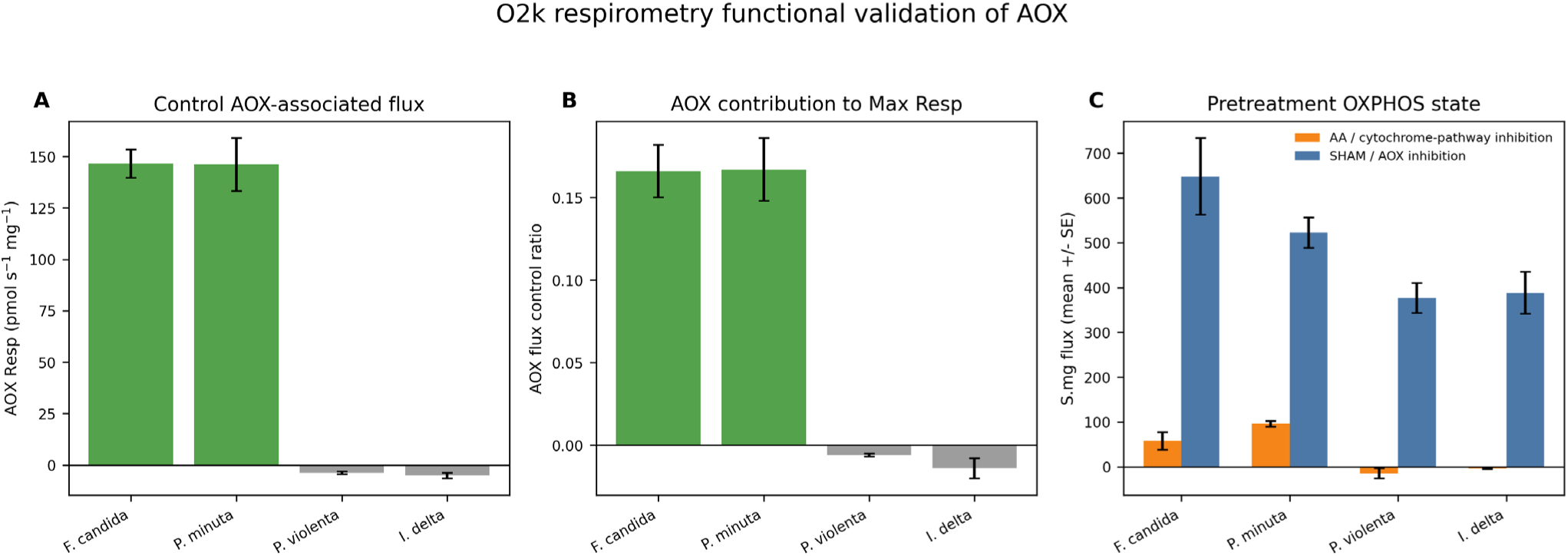
O2k respirometry functional validation of AOX. (A) Control AOX-associated oxygen flux. (B) AOX flux control ratio relative to maximum respiration. (C) Pretreatment S.mg flux after cytochrome-pathway inhibition (AA) or AOX inhibition (SHAM). Bars show means and error bars show SE.

**Figure S7.**
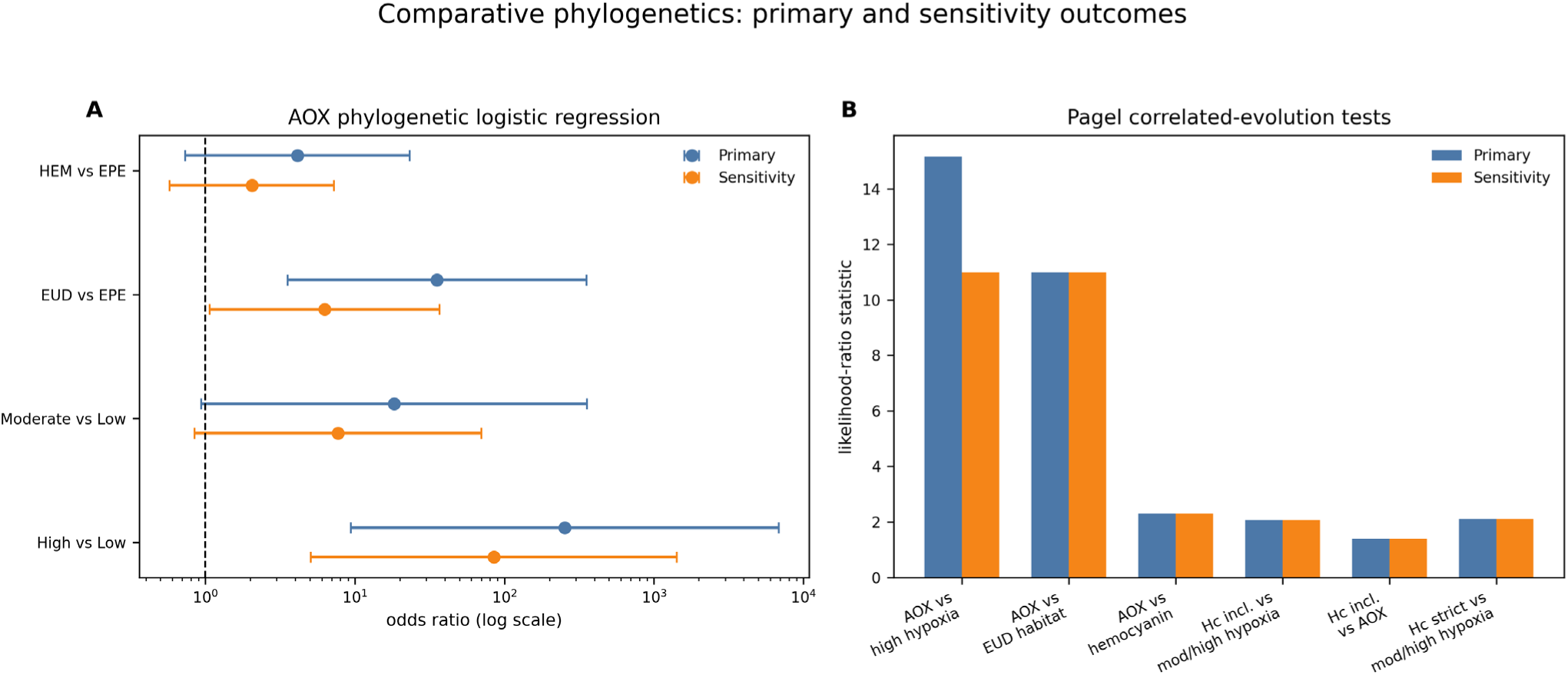
Comparative phylogenetics primary and sensitivity outcomes. (A) Primary and avoidance-aware sensitivity phylogenetic logistic-regression odds ratios for AOX presence. Points show odds ratios and horizontal bars show 95% confidence intervals on a log scale. (B) Primary and sensitivity Pagel likelihood-ratio statistics for AOX and hemocyanin trait-pair tests.

**Figure S8.**
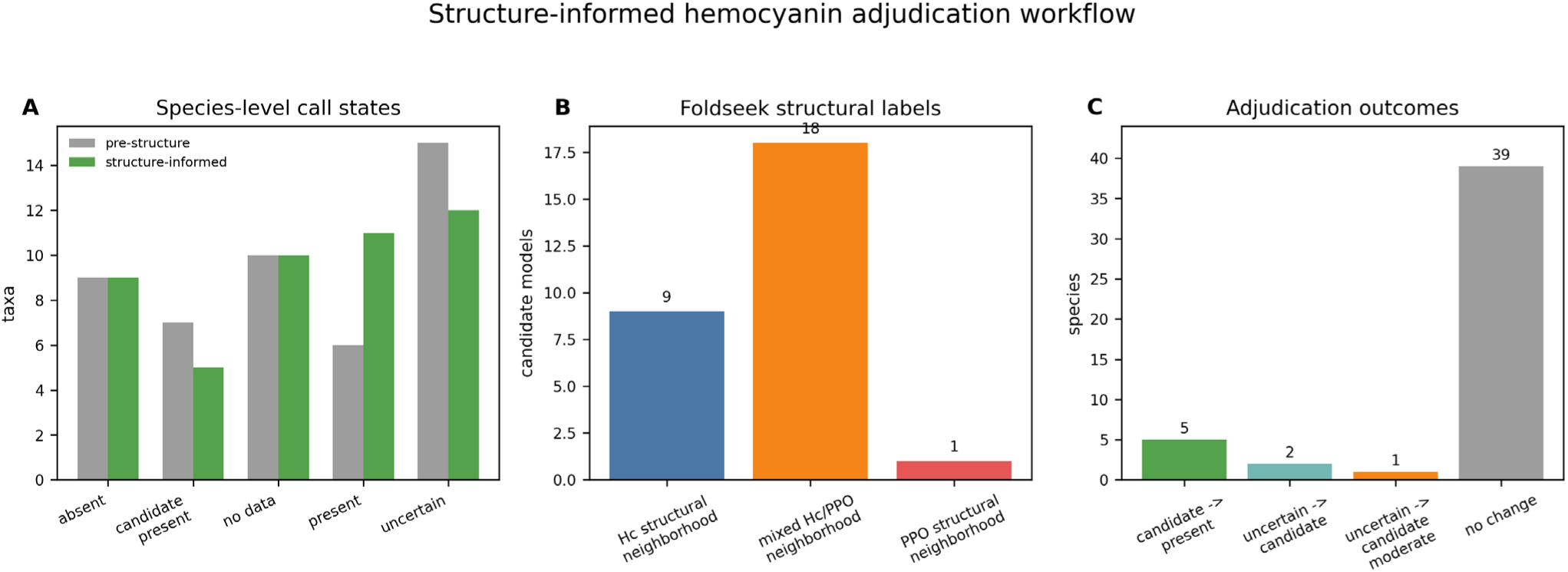
Structure-informed hemocyanin adjudication workflow. (A) Pre-structure and structure-informed species-level call states. (B) Foldseek structural-neighborhood labels among modeled candidate proteins. (C) Final adjudication outcomes, including eight upgrades and no downgrades.

## Tables

**Table S1.**
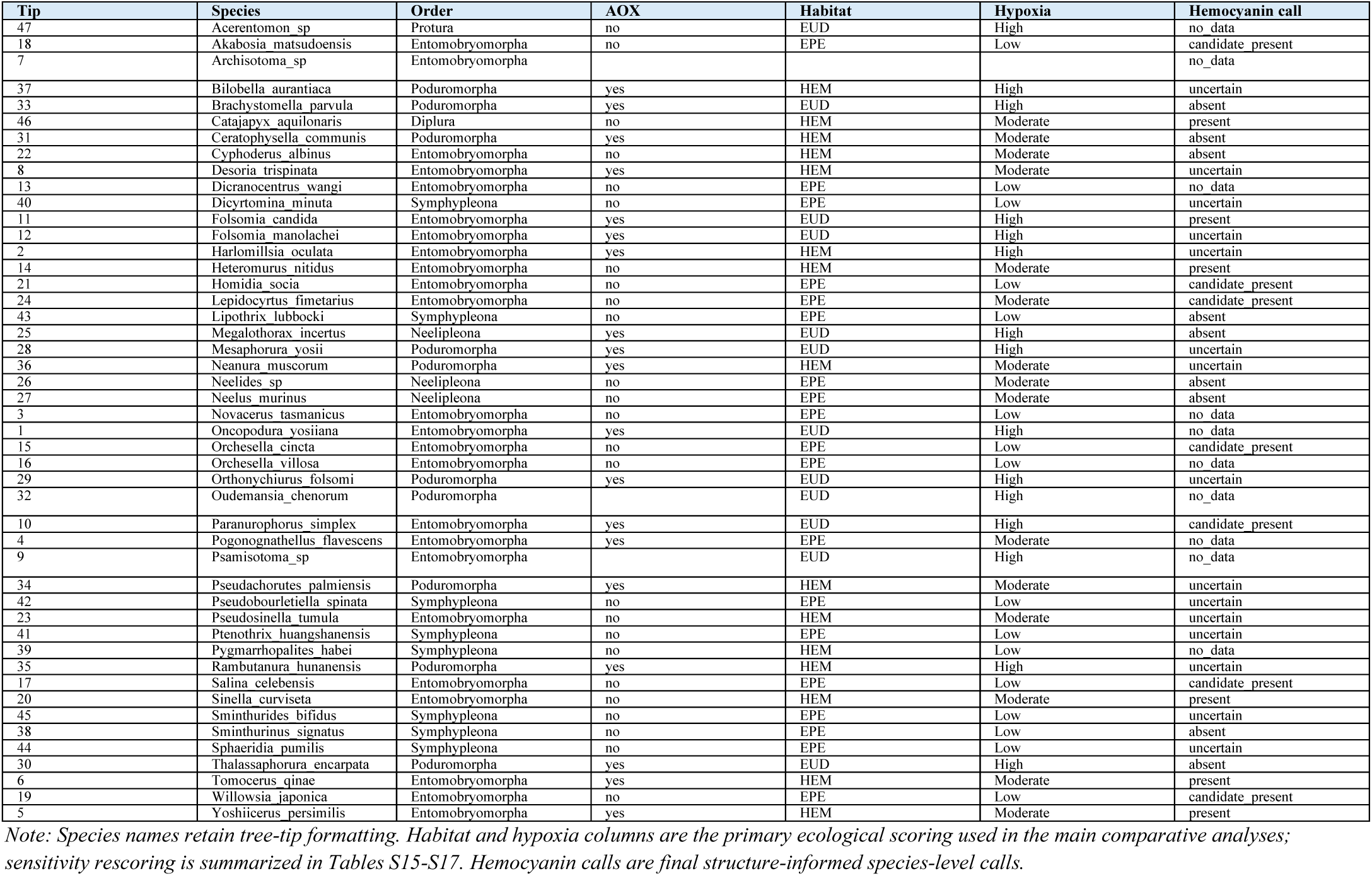
Comparative-phylogenetic trait table for tree tips and outgroups.

**Table S2.** Workflows and data used for appendix.

| Workflow area | Data used |
| --- | --- |
| HGT validation and AOX phylogeny | *F. candida* RefSeq locus validation, read-depth and read-pair checks, rRNA screen, neighborhood taxonomy, TargetP results, AOX phylogeny, and reduced AU topology tests. |
| 202-genome AOX screening/context | Accession-wide miniprot and tblastn scan, context ranking, stitched-candidate confidence, Stage 2B local-context projection, and flanking-interval summaries. |
| Comparative phylogenetics | AOX, habitat, hypoxia, and hemocyanin comparative analyses with sensitivity reruns. |
| Microsynteny | AOX-centered RefSeq marker set, projection to three additional *F. candida* assemblies, sensitivity checks, negative control, and figure exports. |
| O2k respirometry | Curated O2k workflow, respiratory-state mapping, ANOVA/Tukey outputs, FCR calculations, and treatment comparisons. |
| Hemocyanin workflow | Genome-scale hemocyanin/PPO search, HMM refinement, miniprot rescue, ColabFold/Foldseek structure-assisted adjudication, and final call states. |

**Table S3.**
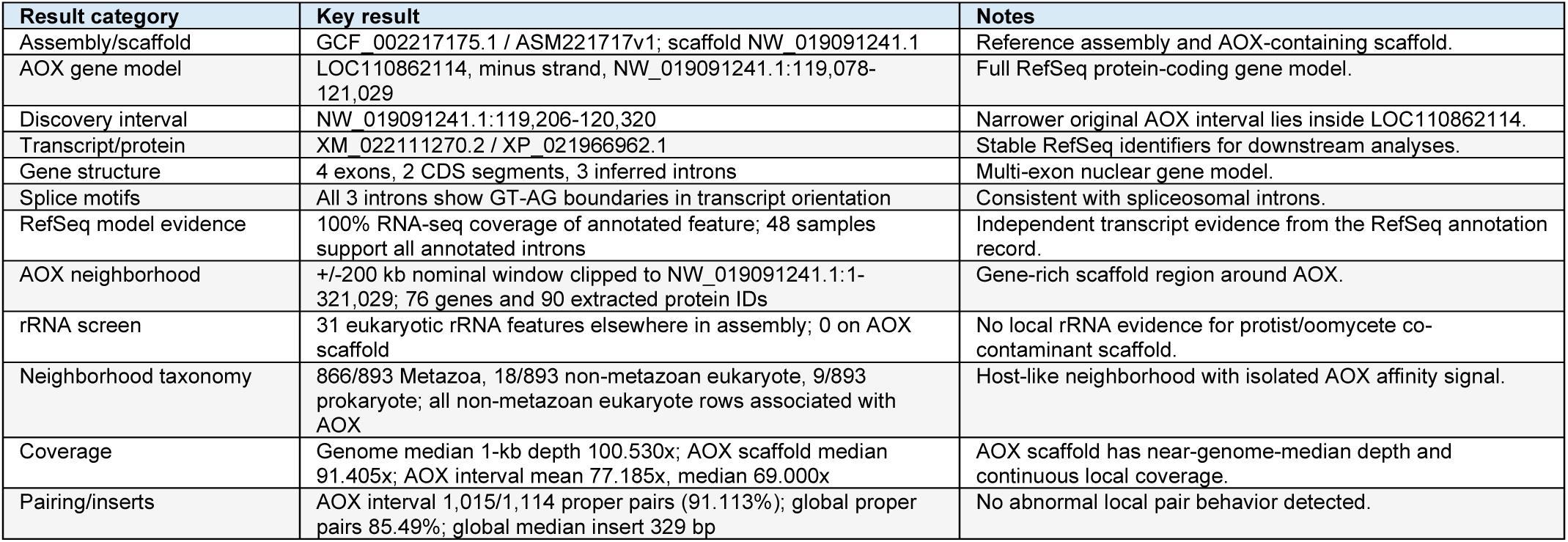
Key *F. candida* AOX locus-validation evidence.

| Result category | Key result | Notes |
| --- | --- | --- |
| Assembly/scaffold | GCF_002217175.1 / ASM221717v1; scaffold NW_019091241.1 | Reference assembly and AOX-containing scaffold. |
| AOX gene model | LOC110862114, minus strand, NW_019091241.1:119,078-121,029 | Full RefSeq protein-coding gene model. |
| Discovery interval | NW_019091241.1:119,206-120,320 | Narrower original AOX interval lies inside LOC110862114. |
| Transcript/protein | XM_022111270.2 / XP_021966962.1 | Stable RefSeq identifiers for downstream analyses. |
| Gene structure | 4 exons, 2 CDS segments, 3 inferred introns | Multi-exon nuclear gene model. |
| Splice motifs | All 3 introns show GT-AG boundaries in transcript orientation | Consistent with spliceosomal introns. |
| RefSeq model evidence | 100% RNA-seq coverage of annotated feature; 48 samples support all annotated introns | Independent transcript evidence from the RefSeq annotation record. |
| AOX neighborhood | +/-200 kb nominal window clipped to NW_019091241.1:1-321,029; 76 genes and 90 extracted protein IDs | Gene-rich scaffold region around AOX. |
| rRNA screen | 31 eukaryotic rRNA features elsewhere in assembly; 0 on AOX scaffold | No local rRNA evidence for protist/oomycete co-contaminant scaffold. |
| Neighborhood taxonomy | 866/893 Metazoa, 18/893 non-metazoan eukaryote, 9/893 prokaryote; all non-metazoan eukaryote rows associated with AOX | Host-like neighborhood with isolated AOX affinity signal. |
| Coverage | Genome median 1-kb depth 100.530x; AOX scaffold median 91.405x; AOX interval mean 77.185x, median 69.000x | AOX scaffold has near-genome-median depth and continuous local coverage. |
| Pairing/inserts | AOX interval 1,015/1,114 proper pairs (91.113%); global proper pairs 85.49%; global median insert 329 bp | No abnormal local pair behavior detected. |

**Table S4.** Splice-junction and iron-binding-site coordinates for *F. candida* AOX.

| Feature | Index or motif | Genomic interval (NW_019091241.1) | Donor / acceptor or aa range | Additional detail |
| --- | --- | --- | --- | --- |
| 1 | Intron 1 | 119,714-119,817 | GT / AG | Pyrimidine % last 20 bases = 60.0 |
| 2 | Intron 2 | 120,330-120,396 | GT / AG | Pyrimidine % last 20 bases = 55.0 |
| 3 | Intron 3 | 120,803-120,887 | GT / AG | Pyrimidine % last 20 bases = 45.0 |
| IBS1 | IBS1 | 119,826-119,852 | 157-165 | CDS/exon segment 119,818-120,329 |
| IBS2 | IBS2 | 119,584-119,613 | 202-211 | CDS/exon segment 119,078-119,713 |
| IBS3 | IBS3 | 119,449-119,484 | 245-256 | CDS/exon segment 119,078-119,713 |
| IBS4 | IBS4 | 119,275-119,304 | 305-314 | CDS/exon segment 119,078-119,713 |

**Table S5.**
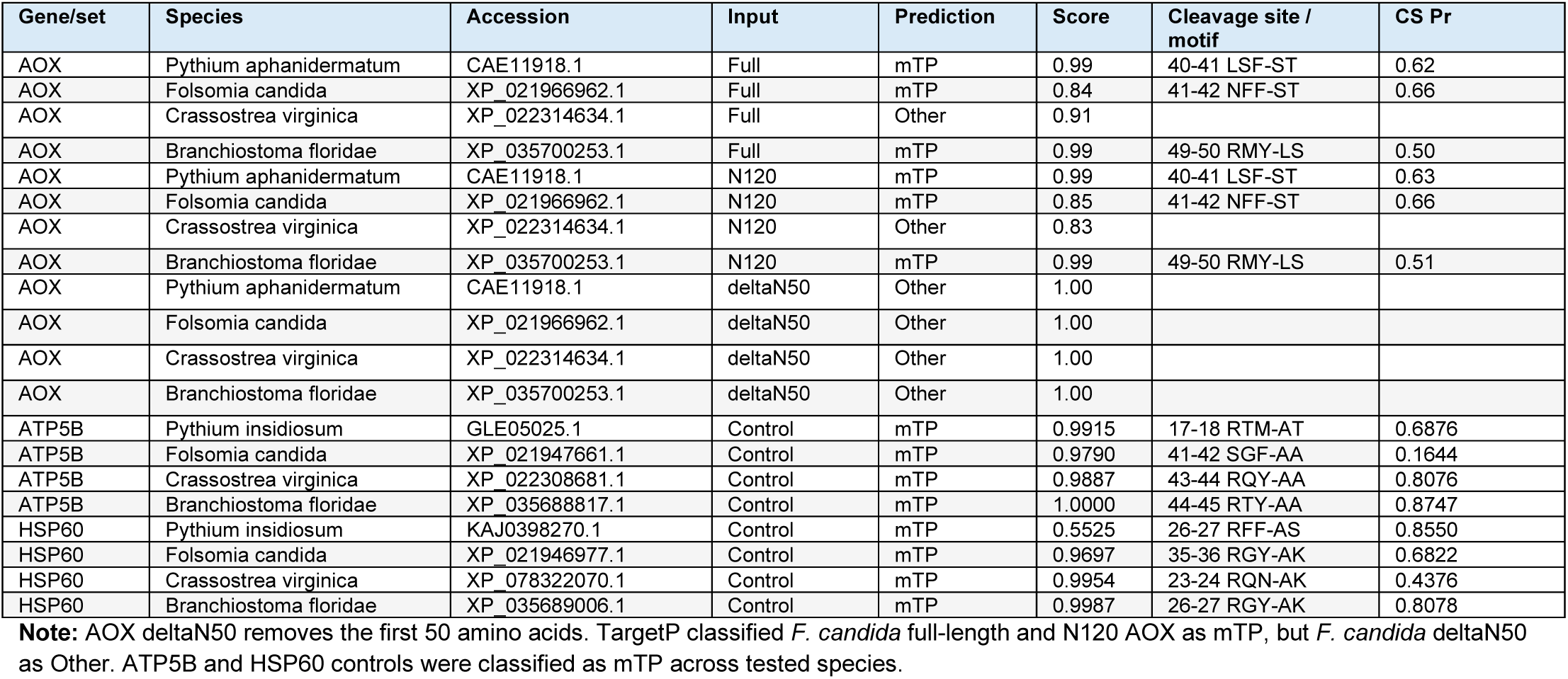
TargetP 2.0 AOX results and mitochondrial positive-control checks.

| Gene/set | Species | Accession | Input | Prediction | Score | Cleavage site / motif | CS Pr |
| --- | --- | --- | --- | --- | --- | --- | --- |
| AOX | *Pythium aphanidermatum* | CAE11918.1 | Full | mTP | 0.99 | 40-41 LSF-ST | 0.62 |
| AOX | *Folsomia candida* | XP_021966962.1 | Full | mTP | 0.84 | 41-42 NFF-ST | 0.66 |
| AOX | *Crassostrea virginica* | XP_022314634.1 | Full | Other | 0.91 |  |  |
| AOX | *Branchiostoma floridae* | XP_035700253.1 | Full | mTP | 0.99 | 49-50 RMY-LS | 0.50 |
| AOX | *Pythium aphanidermatum* | CAE11918.1 | N120 | mTP | 0.99 | 40-41 LSF-ST | 0.63 |
| AOX | *Folsomia candida* | XP_021966962.1 | N120 | mTP | 0.85 | 41-42 NFF-ST | 0.66 |
| AOX | *Crassostrea virginica* | XP_022314634.1 | N120 | Other | 0.83 |  |  |
| AOX | *Branchiostoma floridae* | XP_035700253.1 | N120 | mTP | 0.99 | 49-50 RMY-LS | 0.51 |
| AOX | *Pythium aphanidermatum* | CAE11918.1 | deltaN50 | Other | 1.00 |  |  |
| AOX | *Folsomia candida* | XP_021966962.1 | deltaN50 | Other | 1.00 |  |  |
| AOX | *Crassostrea virginica* | XP_022314634.1 | deltaN50 | Other | 1.00 |  |  |
| AOX | *Branchiostoma floridae* | XP_035700253.1 | deltaN50 | Other | 1.00 |  |  |
| ATP5B | *Pythium insidiosum* | GLE05025.1 | Control | mTP | 0.9915 | 17-18 RTM-AT | 0.6876 |
| ATP5B | *Folsomia candida* | XP_021947661.1 | Control | mTP | 0.9790 | 41-42 SGF-AA | 0.1644 |
| ATP5B | *Crassostrea virginica* | XP_022308681.1 | Control | mTP | 0.9887 | 43-44 RQY-AA | 0.8076 |
| ATP5B | *Branchiostoma floridae* | XP_035688817.1 | Control | mTP | 1.0000 | 44-45 RTY-AA | 0.8747 |
| HSP60 | *Pythium insidiosum* | KAJ0398270.1 | Control | mTP | 0.5525 | 26-27 RFF-AS | 0.8550 |
| HSP60 | *Folsomia candida* | XP_021946977.1 | Control | mTP | 0.9697 | 35-36 RGY-AK | 0.6822 |
| HSP60 | *Crassostrea virginica* | XP_078322070.1 | Control | mTP | 0.9954 | 23-24 RQN-AK | 0.4376 |
| HSP60 | *Branchiostoma floridae* | XP_035689006.1 | Control | mTP | 0.9987 | 26-27 RGY-AK | 0.8078 |

**Note:** AOX deltaN50 removes the first 50 amino acids. TargetP classified *F. candida* full-length and N120 AOX as mTP, but *F. candida* deltaN50 as Other. ATP5B and HSP60 controls were classified as mTP across tested species.

**Table S6.** RefSeq AOX-centered marker set used for *F. candida* microsynteny projection.

| Rank | Side | RefSeq gene | Protein | Reference interval | Strand |
| --- | --- | --- | --- | --- | --- |
| -8 | lower-coordinate flank | LOC110862112 | XP_021966960.1 | NW_019091241.1:70,914-73,118 | - |
| -7 | lower-coordinate flank | LOC110862086 | XP_035701135.1 | NW_019091241.1:74,308-75,828 | - |
| -6 | lower-coordinate flank | LOC110862166 | XP_021967035.1 | NW_019091241.1:81,680-92,446 | - |
| -5 | lower-coordinate flank | LOC110862087 | XP_021966934.2 | NW_019091241.1:96,436-104,886 | + |
| -4 | lower-coordinate flank | LOC110862171 | XP_035701159.1 | NW_019091241.1:104,877-114,104 | - |
| -3 | lower-coordinate flank | LOC110862172 | XP_021967040.1 | NW_019091241.1:106,375-107,563 | + |
| -2 | lower-coordinate flank | LOC110862111 | XP_021966958.1 | NW_019091241.1:114,660-116,942 | + |
| -1 | lower-coordinate flank | LOC110862088 | XP_021966935.1 | NW_019091241.1:116,992-119,070 | + |
| 0 | AOX | LOC110862114 | XP_021966962.1 | NW_019091241.1:119,078-121,029 | - |
| +1 | higher-coordinate flank | LOC110862116 | XP_021966963.1 | NW_019091241.1:121,515-122,928 | - |
| +2 | higher-coordinate flank | LOC110862143 | XP_021967003.1 | NW_019091241.1:124,089-129,176 | - |
| +3 | higher-coordinate flank | LOC110862089 | XP_035701136.1 | NW_019091241.1:130,874-133,044 | - |
| +4 | higher-coordinate flank | LOC118433357 | XP_035701120.1 | NW_019091241.1:133,134-135,551 | - |
| +5 | higher-coordinate flank | LOC110862090 | XP_035701119.1 | NW_019091241.1:136,318-138,325 | - |
| +6 | higher-coordinate flank | LOC110862160 | XP_021967026.1 | NW_019091241.1:139,981-142,453 | + |
| +7 | higher-coordinate flank | LOC110862091 | XP_021966938.2 | NW_019091241.1:142,460-151,905 | - |
| +8 | higher-coordinate flank | LOC110862161 | XP_021967028.1 | NW_019091241.1:154,014-155,559 | + |

**Note:** Side labels are coordinate-side labels relative to AOX, not promoter-direction upstream/downstream for the minus-strand AOX transcript.

**Table S7.** Per-assembly *F. candida* AOX microsynteny support.

| Accession | Assembly level | AOX anchor interval | Anchor source | Target region | Projected/same-side flanks | Support tier |
| --- | --- | --- | --- | --- | --- | --- |
| GCA_020920555.1 | Chromosome | CM037132.1:44,356,368-44,355,850 | aox_tblastn_fragments.tsv | 400,001 bp window | 16/16 | Tier1_strong |
| GCA_052721305.1 | Complete Genome | CP140594.1:45,094,032-45,093,514 | aox_tblastn_fragments.tsv | 400,001 bp window | 16/16 | Tier1_strong |
| GCA_052721315.1 | Complete Genome | CP140601.1:44,923,857-44,923,339 | aox_tblastn_fragments.tsv | 400,001 bp window | 16/16 | Tier1_strong |

**Note:** All target assemblies also passed internal rank-set and all-same-side audits. Window-size and anchor-sensitivity runs remained Tier1\_strong; stitched-anchor sensitivity outputs fell back to fragment anchors in the summarized runs.

**Table S8.**
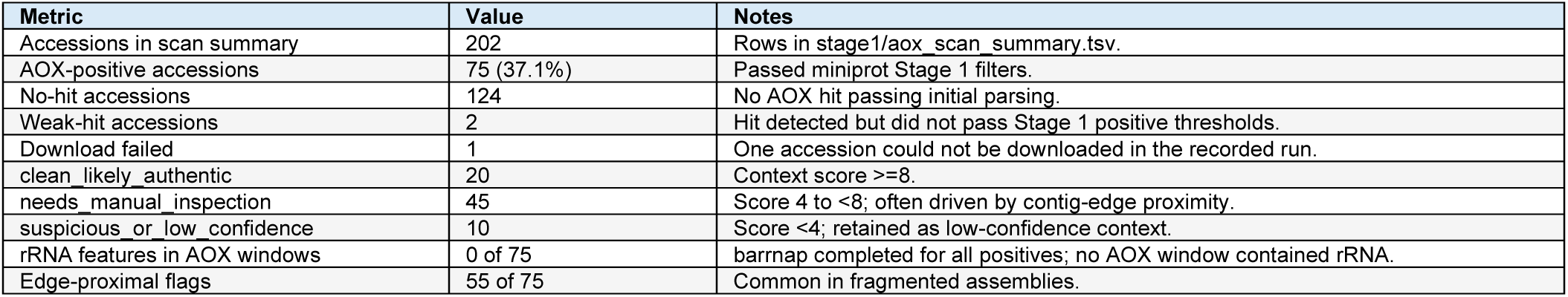
AOX scan and context triage across 202 springtail genome assemblies.

**Table S9.** AOX stitched-candidate and Stage 2B local-context summary.

| Result | Value | Notes |
| --- | --- | --- |
| High-confidence stitched candidate | 65 of 75 (86.7%) | Recovered 3 or 4 of 4 diagnostic AOX iron-binding motifs. |
| Low-confidence stitched set | 10 of 75 | Recovered 1 or 2 motifs. |
| Best candidates with 4 IBS motifs | 50 | Full motif recovery. |
| Best candidates with 3 IBS motifs | 15 | High-confidence by workflow rule. |
| Best candidates with 2 IBS motifs | 8 | Low-confidence retained sequence evidence. |
| Best candidates with 1 IBS motif | 2 | Low-confidence retained sequence evidence. |
| Reference-panel proteins | 88,140 | 37,114 *F. candida* and 51,026 *Allacma fusca* proteins. |
| Raw local projection rows | 12,267 | 5,683 unique reference proteins projected at least once. |
| pass local projections | 6,898 | query fraction $\geq 0.30$, identity $\geq 35\%$, aligned span $\geq 80$ aa. |
| Distinct deduplicated intervals | 608 | 276 left, 255 right, 77 AOX-overlap intervals. |
| Windows with intervals on both sides | 27 of 75 | After family collapse and interval deduplication. |
| Median nearest left/right interval distances | 2,578.5 bp / 2,803.5 bp | Among windows with recovered left or right intervals. |

**Table S10.**
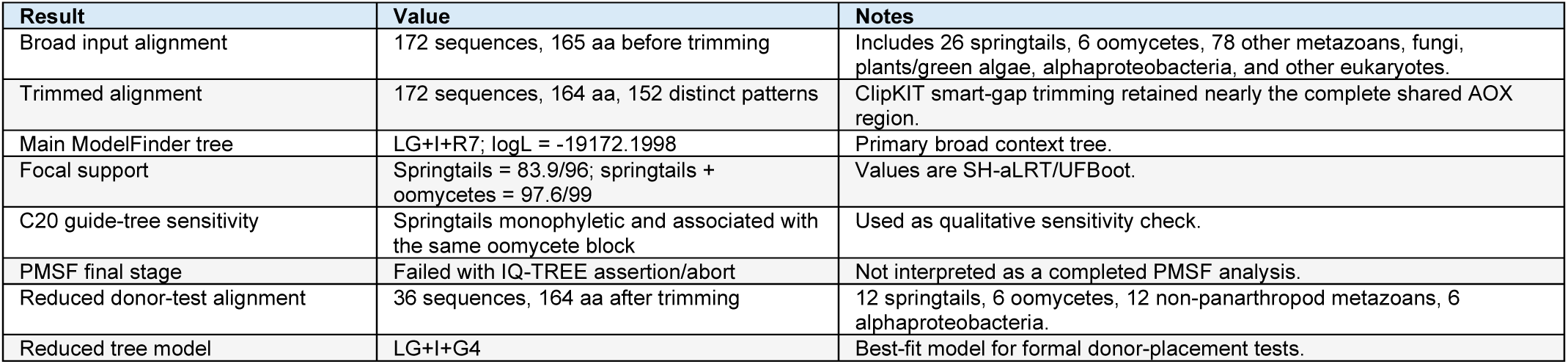
AOX phylogeny workflow and reduced topology-test results.

**Table S11.** Reduced AU topology-test results.

| Tree | Topology | logL | deltaL | p-AU | Status |
| --- | --- | --- | --- | --- | --- |
| 1 | Unconstrained reduced ML | -5091.49978 | 0 | 1 | Not rejected |
| 2 | Oomycota sister | -5091.593981 | 0.0942 | 0.539 | Not rejected |
| 3 | Non-panarthropod metazoa sister | -5109.685389 | 18.186 | 0.00805 | Rejected |

**Table S12.** O2k experimental design, sampling, and respiratory-state mapping.

| Item | Value / variable | Notes |
| --- | --- | --- |
| Active protocol | DLPu / SUIT-008 O2 mt D026 | 2 mL O2k chambers; run-temperature entries approximately |
|  |  | 23-25 C. |
| Titration marks | 1mt, 1PM, 2D, 2c, 3G, 4S, 5U, 7Ama, SHAM, 6Rot | Ten-step user protocol. |
| Species/treatments | *F. candida*: 5 control, 3 AA, 4 SHAM; *P. minuta*: 5, 4, 3; *P. violenta*: 5, 3, 3; *I. delta*: 5, 3, 3 | 46 total observations in curated analysis table. |
| LEAK | PMG.mg | Mass-specific LEAK-linked respiration. |
| CI OXPHOS | G.mg | Complex I OXPHOS state. |
| CI + CII OXPHOS | S.mg | Total OXPHOS state used in pretreatment panel. |
| Max Resp | U.mg | Uncoupled/maximum respiration; denominator for FCR. |
| AOX Resp | AA.mg | AOX-associated oxygen flux after cytochrome-pathway inhibition. |
| Residual | SHAM.mg | Residual flux after SHAM/AOX inhibition. |

**Table S13.** Key O2k respirometry results.

| Result | Value | Notes |
| --- | --- | --- |
| Control AOX Resp ANOVA | F = 141.96, df = 3, p < 0.001 | Strongest state-wise species effect under control conditions. |
| Control AOX FCR ANOVA | F = 62.64, df = 3, p < 0.001 | AOX contribution to maximum respiration differed among species. |
| *F. candida* control AOX Resp | 146.5 +/- 6.9 pmol s<sup>-1</sup> mg<sup>-1</sup>; FCR 0.166 +/- 0.016; n=5 | AOX-positive; HSD group a. |
| *Proisotoma minuta* control AOX Resp | 146.1 +/- 12.8 pmol s<sup>-1</sup> mg<sup>-1</sup>; FCR 0.167 +/- 0.019; n=5 | AOX-positive; HSD group a. |
| *Pseudosinella violenta* control AOX Resp | -3.8 +/- 0.7 pmol s<sup>-1</sup> mg<sup>-1</sup>; FCR -0.006 +/- 0.001; n=5 | AOX-negative; HSD group b. |
| *Isotoma delta* control AOX Resp | -5.1 +/- 1.3 pmol s<sup>-1</sup> mg<sup>-1</sup>; FCR -0.014 +/- 0.006; n=5 | AOX-negative; HSD group b. |
| AA pretreatment S.mg | Species effect F = 23.94, p < 0.001 | AOX-positive species retained positive OXPHOS flux after cytochrome-pathway inhibition. |
| SHAM pretreatment S.mg | Species effect F = 4.54, p = 0.033 | Cytochrome pathway retained when AOX was inhibited. |
| SHAM-treatment AOX Resp state | F = 0.63, p = 0.614 | AOX-specific signal was removed by AOX inhibition. |

**Table S14.**
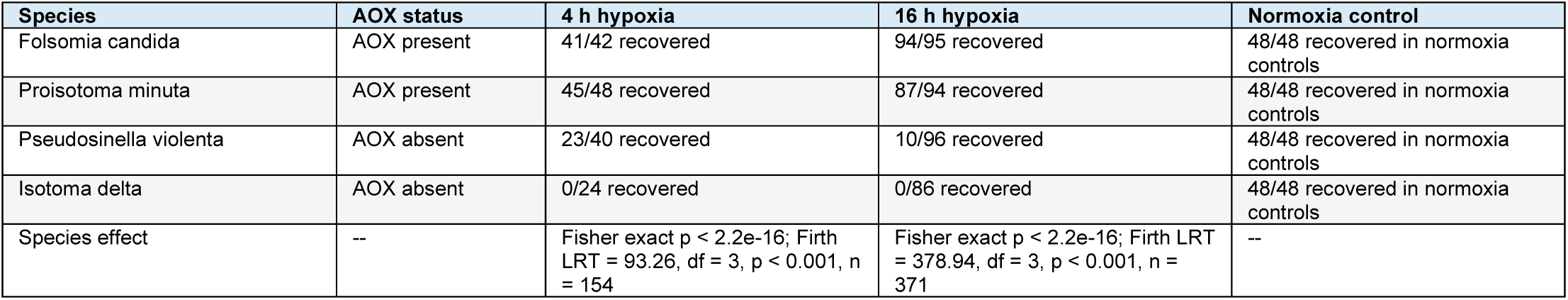
Hypoxia-reoxygenation recovery outcomes.

| Species | AOX status | 4 h hypoxia | 16 h hypoxia | Normoxia control |
| --- | --- | --- | --- | --- |
| *Folsomia candida* | AOX present | 41/42 recovered | 94/95 recovered | 48/48 recovered in normoxia controls |
| *Proisotoma minuta* | AOX present | 45/48 recovered | 87/94 recovered | 48/48 recovered in normoxia controls |
| *Pseudosinella violenta* | AOX absent | 23/40 recovered | 10/96 recovered | 48/48 recovered in normoxia controls |
| *Isotoma delta* | AOX absent | 0/24 recovered | 0/86 recovered | 48/48 recovered in normoxia controls |
| Species effect | -- | Fisher exact p < 2.2e-16; Firth LRT = 93.26, df = 3, p < 0.001, n = 154 | Fisher exact p < 2.2e-16; Firth LRT = 378.94, df = 3, p < 0.001, n = 371 | -- |

**Table S15.** Comparative-phylogenetics dataset scope and AOX trait cross-tabulations.

| Scoring or metric | Category / value | AOX absent | AOX present | Total / notes |
| --- | --- | --- | --- | --- |
| Primary hypoxia | Low | 16 | 0 | 16 |
| Primary hypoxia | Moderate | 7 | 7 | 14 |
| Primary hypoxia | High | 0 | 12 | 12 |
| Sensitivity hypoxia | Low | 14 | 0 | 14 |
| Sensitivity hypoxia | Moderate | 9 | 10 | 19 |
| Sensitivity hypoxia | High | 0 | 9 | 9 |
| Primary habitat | EPE | 18 | 1 | 19 |
| Primary habitat | HEM | 5 | 9 | 14 |
| Primary habitat | EUD | 0 | 9 | 9 |
| Sensitivity habitat | EPE | 15 | 1 | 16 |
| Sensitivity habitat | HEM | 8 | 9 | 17 |
| Sensitivity habitat | EUD | 0 | 9 | 9 |
| Tree tips in ASTRAL backbone | 47 |  |  | All 47 tips match species names in trait tables. |
| AOX-coded taxa in full tree | 44 |  |  | 19 present, 25 absent, 3 missing. |
| AOX regression/Pagel taxa | 42 |  |  | Collembola only after excluding two outgroups; 19 present, 23 absent. |
| Hemocyanin inclusive taxa | 24 |  |  | 15 present and 9 absent among Collembola. |
| Hemocyanin strict taxa | 14 |  |  | 5 present and 9 absent among Collembola. |
| Sensitivity trait changes | 7 |  |  | Seven species have altered habitat and/or hypoxia calls. |

**Table S16.** AOX phylogenetic logistic-regression results under primary and sensitivity ecological scoring.

| Model | Contrast | Primary OR (95% CI); p | Sensitivity OR (95% CI); p | Comparison |
| --- | --- | --- | --- | --- |
| AOX Habitat | HEM vs EPE | 4.121 (0.731-23.22); p = 0.108 | 2.045 (0.577-7.252); p = 0.268 | same direction, not significant |
| AOX Habitat | EUD vs EPE | 35.43 (3.542-354.3); p = 0.0024 | 6.273 (1.072-36.71); p = 0.042 | same direction, significant |
| AOX Hypoxia | Moderate vs Low | 18.25 (0.937-355.3); p = 0.055 | 7.715 (0.848-70.20); p = 0.070 | same direction, marginal |
| AOX Hypoxia | High vs Low | 253.8 (9.408-6847); p = 9.90e-04 | 84.78 (5.054-1422); p = 0.0020 | same direction, significant |

**Table S17.** Pagel correlated-evolution tests under primary and sensitivity scoring.

| Analysis | Primary LR | Primary p | Primary deltaAIC | Sensitivity LR | Sensitivity p | Sensitivity deltaAIC | Comparison |
| --- | --- | --- | --- | --- | --- | --- | --- |
| AOX vs high hypoxia | 15.16 | 5.12e-04 | 11.16 | 10.99 | 0.0041 | 6.990 | same qualitative conclusion |
| AOX vs EUD habitat | 10.99 | 0.0041 | 6.990 | 10.99 | 0.0041 | 6.990 | unchanged |
| AOX vs hemocyanin | 2.307 | 0.315 | -1.693 | 2.307 | 0.315 | -1.693 | unchanged |
| Hemocyanin inclusive vs moderate/high hypoxia | 2.066 | 0.356 | -1.934 | 2.066 | 0.356 | -1.934 | unchanged |
| Hemocyanin inclusive vs AOX | 1.391 | 0.499 | -2.609 | 1.391 | 0.499 | -2.609 | unchanged |
| Hemocyanin strict vs moderate/high hypoxia | 2.113 | 0.715 | -5.887 | 2.113 | 0.715 | -5.887 | unchanged |

**Note:** In the sensitivity workflow, AOX-vs-hypoxia and AOX-vs-habitat Pagel results are identical because High hypoxia and EUD habitat have identical binary vectors after revised scoring.

**Table S18.** Hemocyanin workflow and structure-informed adjudication summary.

| Result | Value | Notes |
| --- | --- | --- |
| Primary taxa targeted by miniprot | 22 | Ambiguous candidate present or uncertain taxa. |
| Non-empty miniprot GFFs | 22 | All targeted taxa produced miniprot output. |
| Translated miniprot proteins | 488 | Rescored against Hc1, Hc2, broad hemocyanin, PPO, and phenoloxidase HMMs. |
| Final unique candidate model sequences | 28 | Submitted to structure-prediction workflow. |
| Reference model sequences | 21 | Hc1, Hc2, broad hemocyanin, and PPO references. |
| ColabFold model jobs | 49 | All 28 candidates and 21 references completed successfully. |
| Foldseek labels | 9 Hc structural, 18 mixed Hc/PPO, 1 PPO structural | Structure used as evidence tier, not stand-alone presence detector. |
| Pre-structure calls | 6 present, 7 candidate present, 15 uncertain, 9 absent, 10 no data | Conservative sequence/HMM/tree layer. |
| Structure-informed calls | 11 present, 5 candidate present, 12 uncertain, 9 absent, 10 no data | Eight upgrades, no downgrades. |

**Table S19.** Species upgraded by structure-informed hemocyanin adjudication.

| Species | Assembly | Prior call | Structure-informed call | Evidence tier |
| --- | --- | --- | --- | --- |
| Akabosia matsudoensis | GCA_019593645.1 | candidate present | present | Tier2 upgrade candidate to present |
| Desoria trispinata | GCA_034695165.1 | uncertain | candidate present | Tier2 upgrade uncertain to candidate |
| Folsomia manolachei | GCA_036491905.1 | uncertain | candidate present | Tier2 upgrade uncertain to candidate |
| Lepidocyrtus fimetarius | GCA_036507315.1 | candidate present | present | Tier2 upgrade candidate to present |
| Orchesella cincta | GCA_034694045.1 | candidate present | present | Tier2 upgrade candidate to present |
| Pseudosinella tumula | GCA_055774745.1 | uncertain | candidate present | Tier2 upgrade uncertain to candidate moderate |
| Salina celebensis | GCA_036561885.1 | candidate present | present | Tier2 upgrade candidate to present |
| Willowsia japonica | GCA_055776465.1 | candidate present | present | Tier2 upgrade candidate to present |

## References

1. K. Christiansen, Behavior and Form in the Evolution of Cave Collembola. Evolution 19, 529–537 (1965).

2. F. G. Howarth, High-Stress Subterranean Habitats and Evolutionary Change in Cave-Inhabiting Arthropods. The American Naturalist 142, S65–S77 (1993).

3. T. L. Poulson, Cave Adaptation in Amblyopsid Fishes. American Midland Naturalist 70, 257 (1963).

4. M. G. Villani, L. L. Allee, A. Díaz, P. S. Robbins, ADAPTIVE STRATEGIES OF EDAPHIC ARTHROPODS. Annu. Rev. Entomol. 44, 233–256 (1999).

5. H. R. Widmer, H. Hoppeler, E. Nevo, C. R. Taylor, E. R. Weibel, Working underground: Respiratory adaptations in the blind mole rat. Proc. Natl. Acad. Sci. U.S.A. 94, 2062–2067 (1997).

6. A. Bundgaard, et al., Low production of mitochondrial reactive oxygen species after anoxia and reoxygenation in turtle hearts. Journal of Experimental Biology 226, jeb245516 (2023).

7. H. M. Honda, P. Korge, J. N. Weiss, Mitochondria and Ischemia/Reperfusion Injury. Annals of the New York Academy of Sciences 1047, 248–258 (2005).

8. T. T. Kozlowski, Soil Aeration, Flooding, and Tree Growth. isa 11, 85–95 (1985).

9. G. Solaini, A. Baracca, G. Lenaz, G. Sgarbi, Hypoxia and mitochondrial oxidative metabolism. Biochimica et Biophysica Acta (BBA) - Bioenergetics 1797, 1171–1177 (2010).

10. A. J. Tompkins, et al., Mitochondrial dysfunction in cardiac ischemia–reperfusion injury: ROS from complex I, without inhibition. Biochimica et Biophysica Acta (BBA) - Molecular Basis of Disease 1762, 223–231 (2006).

11. P. Brookes, V. M. Darley-Usmar, Hypothesis: the mitochondrial NO• signaling pathway, and the transduction of nitrosative to oxidative cell signals: an alternative function for cytochrome C oxidase. Free Radical Biology and Medicine 32, 370–374 (2002).

12. A. Bundgaard, A. M. James, W. Joyce, M. P. Murphy, A. Fago, Suppression of reactive oxygen species generation in heart mitochondria from anoxic turtles: the role of complex I S-nitrosation. Journal of Experimental Biology jeb.174391 (2018). 10.1242/jeb.174391.

13. D. C. Fuhrmann, B. Brüne, Mitochondrial composition and function under the control of hypoxia. Redox Biology 12, 208–215 (2017).

14. P. Gaur, S. Prasad, B. Kumar, S. K. Sharma, P. Vats, High-altitude hypoxia induced reactive oxygen species generation, signaling, and mitigation approaches. Int J Biometeorol 65, 601–615 (2021).

15. I. M. Sokolova, E. P. Sokolov, F. Haider, Mitochondrial Mechanisms Underlying Tolerance to Fluctuating Oxygen Conditions: Lessons from Hypoxia-Tolerant Organisms. Integrative and Comparative Biology 59, 938–952 (2019).

16. H. T. Jacobs, A. L. Moore, There is often – but not always – an alternative! Biochimica et Biophysica Acta (BBA) - Bioenergetics 1866, 149533 (2025).

17. A. E. McDonald, D. V. Gospodaryov, Alternative NAD(P)H dehydrogenase and alternative oxidase: Proposed physiological roles in animals. Mitochondrion 45, 7–17 (2019).

18. R. J. Weaver, A. E. McDonald, Mitochondrial alternative oxidase across the tree of life: Presence, absence, and putative cases of lateral gene transfer. Biochimica et Biophysica Acta (BBA) - Bioenergetics 1864, 149003 (2023).

19. A. Kumari, P. K. Pathak, M. Bulle, A. U. Igamberdiev, K. J. Gupta, Alternative oxidase is an important player in the regulation of nitric oxide levels under normoxic and hypoxic conditions in plants. Journal of Experimental Botany 70, 4345–4354 (2019).

20. J. Selinski, R. Scheibe, D. A. Day, J. Whelan, Alternative Oxidase Is Positive for Plant Performance. Trends in Plant Science 23, 588–597 (2018).

21. G. Vanlerberghe, Alternative Oxidase: A Mitochondrial Respiratory Pathway to Maintain Metabolic and Signaling Homeostasis during Abiotic and Biotic Stress in Plants. IJMS 14, 6805–6847 (2013).

22. R. J. Weaver, Hypothesized Evolutionary Consequences of the Alternative Oxidase (AOX) in Animal Mitochondria. Integrative and Comparative Biology 59, 994–1004 (2019).

23. M. J. Powers, F. S. Barreto, Transcriptomic Adjustment to Decreasing Oxygen Reveals Novel Functional Strategies for Extreme Hypoxia Tolerance in the Copepod Tigriopus californicus. Genome Biology and Evolution 18 (2026).

24. A. Andjelković, et al., Diiron centre mutations in Ciona intestinalis alternative oxidase abolish enzymatic activity and prevent rescue of cytochrome oxidase deficiency in flies. Sci Rep 5, 18295 (2015).

25. R. El-Khoury, M. Rak, P. Bénit, H. T. Jacobs, P. Rustin, Cyanide resistant respiration and the alternative oxidase pathway: A journey from plants to mammals. Biochimica et Biophysica Acta (BBA) - Bioenergetics 1863, 148567 (2022).

26. O. A. Borovik, O. I. Grabelnych, Mitochondrial alternative cyanide-resistant oxidase is involved in an increase of heat stress tolerance in spring wheat. Journal of Plant Physiology 231, 310–317 (2018).

27. R. Clifton, A. H. Millar, J. Whelan, Alternative oxidases in Arabidopsis: A comparative analysis of differential expression in the gene family provides new insights into function of non-phosphorylating bypasses. Biochimica et Biophysica Acta (BBA) - Bioenergetics 1757, 730–741 (2006).

28. L. Duvenage, et al., Alternative oxidase induction protects Candida albicans from respiratory stress and promotes hyphal growth. [Preprint] (2018). Available at: http://biorxiv.org/lookup/doi/10.1101/405670 [Accessed 27 March 2026].

29. E. Sierra-Campos, I. Velázquez, D. Matuz-Mares, A. Villavicencio-Queijeiro, J. P. Pardo, Functional properties of the Ustilago maydis alternative oxidase under oxidative stress conditions. Mitochondrion 9, 96–102 (2009).

30. F. E. Sluse, W. Jarmuszkiewicz, Alternative oxidase in the branched mitochondrial respiratory network: an overview on structure, function, regulation, and role. Braz J Med Biol Res 31, 733–747 (1998).

31. R. Sussarellu, et al., Rapid mitochondrial adjustments in response to short-term hypoxia and re-oxygenation in the Pacific oyster *Crassostrea gigas*. Journal of Experimental Biology jeb.075879 (2013). 10.1242/jeb.075879.

32. O. Blokhina, K. V. Fagerstedt, Oxidative metabolism, ROS and NO under oxygen deprivation. Plant Physiology and Biochemistry 48, 359–373 (2010).

33. R. Chen, et al., Reactive Oxygen Species Formation in the Brain at Different Oxygen Levels: The Role of Hypoxia Inducible Factors. Front. Cell Dev. Biol. 6, 132 (2018).

34. T. L. Clanton, Hypoxia-induced reactive oxygen species formation in skeletal muscle. Journal of Applied Physiology 102, 2379–2388 (2007).

35. M. Erecińska, I. A. Silver, Tissue oxygen tension and brain sensitivity to hypoxia. Respiration Physiology 128, 263–276 (2001).

36. A. E. McDonald, G. C. Vanlerberghe, J. F. Staples, Alternative oxidase in animals: unique characteristics and taxonomic distribution. Journal of Experimental Biology 212, 2627–2634 (2009).

37. H. Gisin, Okologie und levensgemenischaften der collembolen im schweizerischen exkursionsgebiet basels. Revue suisse de Zoologie 50, 131–224 (1943).

38. C. Li, et al., Euedaphic Rather than Hemiedaphic or Epedaphic Collembola Are More Sensitive to Different Climate Conditions in the Black Soil Region of Northeast China. Insects 16, 275 (2025).

39. A. A. Potapov, E. E. Semenina, A. Yu. Korotkevich, N. A. Kuznetsova, A. V. Tiunov, Connecting taxonomy and ecology: Trophic niches of collembolans as related to taxonomic identity and life forms. Soil Biology and Biochemistry 101, 20–31 (2016).

40. E. Nazari, S. Besharat, K. Zeinalzadeh, A. Mohammadi, Measurement and simulation of the water flow and root uptake in soil under subsurface drip irrigation of apple tree. Agricultural Water Management 255, 106972 (2021).

41. J. Neira, M. Ortiz, L. Morales, E. Acevedo, Oxygen diffusion in soils: Understanding the factors and processes needed for modeling. *Chilean J*. Agric. Res. 75, 35–44 (2015).

42. S. Flachsbarth, M. Kruse, T. Burmester, Distribution and hypoxia-regulation of haemocyanin in springtails (Collembola). Insect Molecular Biology 26, 633–641 (2017).

43. R. El-Khoury, et al., Alternative Oxidase Expression in the Mouse Enables Bypassing Cytochrome c Oxidase Blockade and Limits Mitochondrial ROS Overproduction. PLoS Genet 9, e1003182 (2013).

44. D. J. M. Fernandez-Ayala, et al., Expression of the Ciona intestinalis Alternative Oxidase (AOX) in Drosophila Complements Defects in Mitochondrial Oxidative Phosphorylation. Cell Metabolism 9, 449–460 (2009).

45. G. A. J. Hakkaart, E. P. Dassa, H. T. Jacobs, P. Rustin, Allotopic expression of a mitochondrial alternative oxidase confers cyanide resistance to human cell respiration. EMBO Rep 7, 341–345 (2006).

46. A. E. McDonald, S. M. Sieger, G. C. Vanlerberghe, Methods and approaches to study plant mitochondrial alternative oxidase. Physiologia Plantarum 116, 135–143 (2002).

47. B. A. P. Williams, et al., A Broad Distribution of the Alternative Oxidase in Microsporidian Parasites. PLoS Pathog 6, e1000761 (2010).

48. J. Jayawardhane, et al., Roles for Plant Mitochondrial Alternative Oxidase Under Normoxia, Hypoxia, and Reoxygenation Conditions. Front. Plant Sci. 11, 566 (2020).

49. D. Cantoni, et al., Localization and functional characterization of the alternative oxidase in Naegleria. J Eukaryotic Microbiology 69, e12908 (2022).

50. S. Völkel, M. K. Grieshaber, Mitochondrial Sulfide Oxidation in *Arenicola Marina*: Evidence for Alternative Electron Pathways. European Journal of Biochemistry 235, 231–237 (1996).

51. K. Bremer, H. Yasuo, P. V. Debes, H. T. Jacobs, The alternative oxidase (AOX) increases sulphide tolerance in the highly invasive marine invertebrate *Ciona intestinalis*. Journal of Experimental Biology 224, jeb242985 (2021).

52. Z. Liu, et al., Multifactor transcriptional control of alternative oxidase induction integrates diverse environmental inputs to enable fungal virulence. Nat Commun 14, 4528 (2023).

53. B. Szal, Y. Jolivet, M. Hasenfratz-Sauder, P. Dizengremel, A. M. Rychter, Oxygen concentration regulates alternative oxidase expression in barley roots during hypoxia and post-hypoxia. Physiologia Plantarum 119, 494–502 (2003).

54. M. S. Yusseppone, et al., Inducing the Alternative Oxidase Forms Part of the Molecular Strategy of Anoxic Survival in Freshwater Bivalves. Front. Physiol. 9, 100 (2018).

55. A. Fago, F. B. Jensen, Hypoxia Tolerance, Nitric Oxide, and Nitrite: Lessons From Extreme Animals. Physiology 30, 116–126 (2015).

56. M. Hermes-Lima, et al., Preparation for oxidative stress under hypoxia and metabolic depression: Revisiting the proposal two decades later. Free Radical Biology and Medicine 89, 1122–1143 (2015).

57. J. Jayawardhane, et al., Roles for Plant Mitochondrial Alternative Oxidase Under Normoxia, Hypoxia, and Reoxygenation Conditions. Front. Plant Sci. 11, 566 (2020).

58. V. I. Lushchak, Environmentally induced oxidative stress in aquatic animals. Aquatic Toxicology 101, 13–30 (2011).

59. K. E. Van Holde, K. I. Miller, H. Decker, Hemocyanins and Invertebrate Evolution. Journal of Biological Chemistry 276, 15563–15566 (2001).

60. N. Monesi, et al., Identification and characterization of a laterally transferred alternative oxidase (AOX) in a terrestrial insect, the dipteran Pseudolycoriella hygida. Biochimie 233, 60–74 (2025).

61. H. Li, Protein-to-genome alignment with miniprot. Bioinformatics 39, btad014 (2023).

62. J. L. Steenwyk, T. J. Buida, Y. Li, X.-X. Shen, A. Rokas, ClipKIT: A multiple sequence alignment trimming software for accurate phylogenomic inference. PLoS Biol 18, e3001007 (2020).

63. B. Q. Minh, et al., IQ-TREE 2: New Models and Efficient Methods for Phylogenetic Inference in the Genomic Era. Molecular Biology and Evolution 37, 1530–1534 (2020).

64. A. Potapov, et al., Towards a global synthesis of Collembola knowledge – challenges and potential solutions. Soil Organisms 92, 161–188 (2020).

65. L. J. Revell, phytools 2.0: an updated R ecosystem for phylogenetic comparative methods (and other things). PeerJ 12, e16505 (2024).

66. L. S. Tung Ho, C. Ané, A Linear-Time Algorithm for Gaussian and Non-Gaussian Trait Evolution Models. Systematic Biology 63, 397–408 (2014).

## SI References

1. P. Danecek, et al., Twelve years of SAMtools and BCFtools. GigaScience 10, giab008 (2021).

2. O. Emanuelsson, H. Nielsen, S. Brunak, G. Von Heijne, Predicting Subcellular Localization of Proteins Based on their N-terminal Amino Acid Sequence. J. Mol. Biol. 300, 1005–1016 (2000).

3. H. Li, Protein-to-genome alignment with miniprot. Bioinformatics 39, btad014 (2023).

4. R. J. Weaver, A. E. McDonald, Mitochondrial alternative oxidase across the tree of life: Presence, absence, and putative cases of lateral gene transfer. Biochim. Biophys. Acta BBA - Bioenerg. 1864, 149003 (2023).

5. J. L. Steenwyk, T. J. Buida, Y. Li, X.-X. Shen, A. Rokas, ClipKIT: A multiple sequence alignment trimming software for accurate phylogenomic inference. PLOS Biol. 18, e3001007 (2020).

6. T. K. Wong, et al., IQ-TREE 3: phylogenomic inference software using complex evolutionary models. [Preprint] (2025).

7. S. Kalyaanamoorthy, B. Q. Minh, T. K. F. Wong, A. Von Haeseler, L. S. Jermiin, ModelFinder: fast model selection for accurate phylogenetic estimates. Nat. Methods 14, 587–589 (2017).

8. D. T. Hoang, O. Chernomor, A. Von Haeseler, B. Q. Minh, L. S. Vinh, UFBoot2: Improving the Ultrafast Bootstrap Approximation. Mol. Biol. Evol. 35, 518–522 (2018).

9. D. Yu, et al., Whole-genome-based phylogenetic analyses provide new insights into the evolution of springtails (Hexapoda: Collembola). Mol. Phylogenet. Evol. 200, 108169 (2024).

10. A. A. Potapov, E. E. Semenina, A. Yu. Korotkevich, N. A. Kuznetsova, A. V. Tiunov, Connecting taxonomy and ecology: Trophic niches of collembolans as related to taxonomic identity and life forms. Soil Biol. Biochem. 101, 20–31 (2016).

11. C. Li, et al., Euedaphic Rather than Hemiedaphic or Epedaphic Collembola Are More Sensitive to Different Climate Conditions in the Black Soil Region of Northeast China. Insects 16, 275 (2025).

12. A. Potapov, et al., Towards a global synthesis of Collembola knowledge – challenges and potential solutions. Soil Org. 92, 161–188 (2020).

13. T. T. Kozlowski, Soil Aeration, Flooding, and Tree Growth. Arboric. Urban For. 11, 85–95 (1985).

14. J. Neira, M. Ortiz, L. Morales, E. Acevedo, Oxygen diffusion in soils: Understanding the factors and processes needed for modeling. *Chil*. J. Agric. Res. 75, 35–44 (2015).

15. W. L. Silver, A. E. Lugo, M. Keller, Soil oxygen availability and biogeochemistry along rainfall and topographic gradients in upland wet tropical forest soils. Biogeochemistry 44, 301–328 (1999).

16. M. Bollazzi, D. Römer, F. Roces, Carbon dioxide levels and ventilation in *Acromyrmex* nests: significance and evolution of architectural innovations in leaf-cutting ants. R. Soc. Open Sci. 8, 210907 (2021).

17. C. Kleineidam, F. Roces, Carbon dioxide concentrations and nest ventilation in nests of the leaf-cutting ant Atta vollenweideri: Insectes Sociaux 47, 241–248 (2000).

18. S. R. Eddy, Accelerated Profile HMM Searches. PLoS Comput. Biol. 7, e1002195 (2011).

19. M. Mirdita, et al., ColabFold: making protein folding accessible to all. Nat. Methods 19, 679–682 (2022).

20. M. Van Kempen, et al., Fast and accurate protein structure search with Foldseek. Nat. Biotechnol. 42, 243–246 (2024).

21. L. J. Revell, phytools 2.0: an updated R ecosystem for phylogenetic comparative methods (and other things). PeerJ 12, e16505 (2024).

22. L. S. Tung Ho, C. Ané, A Linear-Time Algorithm for Gaussian and Non-Gaussian Trait Evolution Models. Syst. Biol. 63, 397–408 (2014).

